# Mutations in the Insulator Protein Suppressor of Hairy Wing Induce Genome Instability

**DOI:** 10.1101/551002

**Authors:** Shih-Jui Hsu, Emily C. Stow, James R. Simmons, Heather A. Wallace, Andrea Mancheno Lopez, Shannon Stroud, Mariano Labrador

**Affiliations:** Department of Biochemistry and Cellular and Molecular Biology, The University of Tennessee, Knoxville, TN 37996, USA

**Keywords:** Chromatin insulators, chromatin boundaries, gypsy, suppressor of Hairy wing, Su(Hw), Drosophila, oogenesis, Gurken, ATR, ATR mediated DNA damage response, MTOC, Double strand breaks (DSBs), γH2Av, H4K20me1, Chk1, grapes, endoreplication, replication stress

## Abstract

Chromatin insulator proteins mediate the formation of contacts between distant insulator sites along chromatin fibers. Long-range contacts facilitate communication between regulatory sequences and gene promoters throughout the genome, allowing accurate gene transcription regulation during embryo development and cell differentiation. Lack of insulator function has detrimental effects often resulting in lethality. The *Drosophila* insulator protein Suppressor of Hairy wing [Su(Hw)] is not essential for viability, but plays a crucial role in female oogenesis. The mechanism(s) by which Su(Hw) promotes proper oogenesis remains unclear. To gain insight into the functional properties of chromatin insulators, we further characterize the oogenesis phenotypes of *su(Hw)* mutant females. We find that mutant egg chambers frequently display an irregular number of nurse cells, have poorly formed microtubule organization centers (MTOC) in the germarium, and show mislocalized Gurken (Grk) in later stages of oogenesis. Furthermore, eggshells produced by partially rescued *su(Hw)* mutant females exhibit dorsoventral patterning defects that are identical to defects found in spindle mutants or in piRNA pathway mutants. Further analysis reveals an excess of DNA damage in egg chambers, which is independent of activation of transposable elements, and that Gurken localization defects and oogenesis progression are partially rescued by mutations in *mei-41* and *chk1* genes. In addition, we show that Su(Hw) is required for chromosome integrity in dividing neuroblasts from larval brains. Together, these findings suggest that Su(Hw) plays a critical role in maintaining genome integrity during germline development in *Drosophila* females as well as in dividing somatic cells.

## Introduction

Chromatin insulators facilitate higher-order chromatin organization in the nucleus by stabilizing interactions between distant sites in the chromatin fiber. These long-range contacts help orchestrate interactions between regulatory sequences and gene promoters to accommodate the complex genomic networks of gene transcription required to promote cell and tissue differentiation during embryo development (Labrador and Corces, 2002; Lupianez et al., 2015; Yang and Corces, 2012). Insulator function is conserved throughout eukaryotes (Gurudatta and Corces, 2009; Heger and Wiehe, 2014; Schoborg and Labrador, 2014; Schoborg and Labrador, 2010; Van Bortle and Corces, 2012b). Canonical insulator properties include the ability to prevent communication between distal enhancers and promoters when positioned in between them and function as boundaries to protect genes against heterochromatin-mediated silencing (Brasset and Vaury, 2005; Gaszner and Felsenfeld, 2006; Kellum and Schedl, 1991; Roseman et al., 1993; Udvardy et al., 1985; West et al., 2002; Zhao et al., 1995). These properties are mediated by insulator proteins, which bind the insulator DNA and may facilitate long-range DNA-DNA interactions (Phillips-Cremins and Corces, 2013; Van Bortle and Corces, 2012a). Recent progress in the study of insulator protein distribution and in the analysis of the three-dimensional organization of the genome within the nucleus has revealed a role of these proteins in the architectural organization of the genome (Lieberman-Aiden et al., 2009; Negre et al., 2010; Ong and Corces, 2014; Phillips-Cremins and Corces, 2013; Rao et al., 2014; Rowley et al., 2017). Thus, a new paradigm has emerged where insulator proteins in combination with cohesin establish topologically associating domains of DNA, which correspond with topologically independent regions of gene transcription activation or repression throught the genome (Fudenberg et al., 2016; Smith et al., 2016). However, although the functional principles behind this organization are strongly supported by experimental ovservations in mammalian cells, the question of whether this model of genome organization is universal amongst euckayotes, as well as the implications of the model in genome organization, genome function and genome integrity, remain an active focus of research (Canela et al., 2017; Oomen et al., 2019; Rowley et al., 2017).

The *gypsy* retrotransposon found in *Drosophila* contains one of the earliest characterized insulators. *Gypsy* can integrate at sites between enhancers and promoters, thereby disrupting enhancer-promoter communication and causing mutations that can be suppressed by mutations in the *suppressor of Hairy wing* gene [*su(Hw)*] (Modolell et al., 1983; Parkhurst and Corces, 1986; Spana et al., 1988). In addition to Su(Hw), which directly binds to the insulator DNA, two other proteins are required for *gypsy* insulator function: Modifier of mdg4 [Mod(mdg4)-67.2] and Centrosomal Protein 190 (CP190), which directly interact with Su(Hw) (Georgiev and Kozycina, 1996; Georgiev and Gerasimova, 1989; Ghosh et al., 2001; Pai et al., 2004). Unlike other insulator proteins in *Drosophila* such as dCTCF, CP190, BEAF, or GAGA factor, the function of both Su(Hw) and its binding partner Mod(mdg4)-67.2 are dispensable for viability (Butcher et al., 2004; Gerasimova et al., 1995; Katokhin et al., 2001; Klug et al., 1968; Mohan et al., 2007; Roy et al., 2007). CP190 has insulator activity that is independent from Su(Hw) and forms insulators in the genome in association with other insulator proteins (Bushey et al., 2009; Mohan et al., 2007; Moshkovich et al., 2011). Homozygous *su(Hw)* loss-of-function mutations are viable with no evident phenotype other than female sterility (Klug et al., 1968; Klug et al., 1970). In ovaries, Su(Hw) is detected in the nucleus of both somatic follicle cells and germ cells (Baxley et al., 2011). Specifically, loss of Su(Hw) leads to suppression of yolk deposition in the oocyte and oocyte development is arrested at mid-oogenesis (Harrison et al., 1993; Klug et al., 1968; Klug et al., 1970). More recent findings have revealed that loss of Su(Hw) leads to an upregulation of neuronal gene expression in germline tissue, suggesting that missexpression of these genes could be responsible for the sterility phenotype of *su(Hw)* mutations (Baxley et al., 2011; Harrison et al., 1993; Soshnev et al., 2013; Soshnev et al., 2012). In fact, the oogenesis phenotype in *su(Hw)* mutant females can be partially suppressed by mutations that reduce the expression of the RNA-binding protein 9 (Rbp9), a protein expressed at higher levels in *su(Hw)* mutant ovaries that is involved in blood-brain barrier establishment (Kim et al., 2010; Soshnev et al., 2013). Rescued females, however, do not produce viable offspring, and eggshells from laid embryos reveal strong dorsoventral transformations. Given the complexity of this phenotype, analysis of *su(Hw)* mutations in the female germ line could be instrumental for further understanding the role of Su(Hw) in oogenesis and the function of insulator proteins during development in general.

In *Drosophila*, oogenesis begins with the asymmetric cell division of a germline stem cell located at the tip of the germarium, which gives rise to a daughter stem cell and a cystoblast. The cystoblast undergoes four incomplete mitotic divisions, forming an egg chamber containing sixteen germ cells that remain interconnected and are enclosed by an epithelium of follicle cells. Only one of the sixteen germ cells will adopt the oocyte cell fate while the remaining fifteen cells become nurse cells. Each egg chamber undergoes a developmental process that culminates with the formation of a mature oocyte at stage 14. As oogenesis progresses, at stage 6, the nucleus of nurse cells undergoes a dramatic change from a condensed five-blobs chromosome configuration to a decondensed chromosome morphology (Bate and Martinez Arias, 1993; Dej and Spradling, 1999). Before mid-oogenesis arrest, the only visible chromatin-configuration defect in *su(Hw)* mutant egg chambers is a delayed chromatin dispersal of nurse cell polytene chromosomes at stage 7 or 8. The prolonged development of defective egg chambers is eliminated by mid-oogenesis arrest resulting in egg chamber degeneration around stages 9 to 10 (Baxley et al., 2011; Harrison et al., 1993; Klug et al., 1968). This defective chromatin dispersal is a common trait among a large number of unrelated mutants, such as genes encoding the splicesosome component *prp22* (Klusza et al., 2013), the piwi-interacting RNAs (piRNAs) related protein, and *rhino* (Klattenhoff et al., 2009; Volpe et al., 2001), which complicates the identification of the mechanisms associated with Su(Hw) activity in the genome of egg chamber cells.

In addition to defects resulting from mutations in oogenesis genes, uncontrolled transposon activity may also cause a severe disruption of development during oogenesis. piRNAs are critical molecules involved in suppression of transposable element activity. These RNA sequences were originally called repeat-associated siRNAs (rasi-RNAs) as they are derived from repetitive sequences in the genome (Aravin et al., 2003; Brennecke et al., 2007; Feschotte, 2008; Yin and Lin, 2007). Mutations in genes involved in piRNA production such as *aubergine* (*aub*), *spindle-E* (*spnE*), and *maelstrom* (*mael*) result in transposable element overexpression and mobilization that create double strand breaks (DSBs) in the host genome (Chen et al., 2007; Cook et al., 2004; Klattenhoff et al., 2007; Sienski et al., 2012). The unleashed retrotransposon activation triggers the DNA damage response, causing defects in microtubule polarization in early oogenesis that further disrupt Gurken signaling in later stages that are essential for dorsoventral determination in embryonic development (Chen et al., 2007; Khurana and Theurkauf, 2010; Klattenhoff et al., 2007). Interestingly, the DNA damage response resulting from over-activation of retrotransposons has been known to utilize the same pathway triggered by unrepaired DSBs in mutants of spindle class genes encoding meiotic DNA damage repair proteins. These unrepaired DSBs activate meiotic checkpoints mediated by *mei-41* (ataxia telangiectasia-related-ATR-ortholog) and *mnk* (checkpoint kinase 2-chk2-ortholog) (Abdu et al., 2002; Ghabrial and Schupbach, 1999; LaRocque et al., 2007; Staeva-Vieira et al., 2003).

In this study, we conclude that the loss of Su(Hw) creates an accumulation of non-meiotic DNA damage in germline cells of ovaries, thereby activating DNA damage checkpoints. We show that DNA damage is independent of transposable element activity and that mutations in meiotic checkpoint genes *mei-41* (ATR) and *grapes* (*chk1*), but not *mnk* (*chk2, loki*), result in the rescue of the *su(Hw)* mutant spindle phenotype in ovaries. We conclude that the lack of *su(Hw)* expression in ovaries provokes an excess of unrepaired DNA breaks in germline cells, and propose that this DNA damage is caused by replication stress. Our data supports that replication stress occurs in dividing somatic cells as well, in *su(Hw* mutants.

## Results

### Egg chambers from *su(Hw)* mutant ovaries exhibit an irregular number of nurse cells

Loss of Su(Hw) causes incomplete oocyte development, resulting in egg chamber degeneration at mid-oogenesis (Baxley et al., 2011; Harrison et al., 1993; Klug et al., 1968; Klug et al., 1970). Previous observations identified an increased number of nurse cells in a fraction of egg chambers from homozygous *su(Hw)*^*v*^ and *su(Hw)*^*e04061/2*^ trans-heterozygous mutants, but this phenotype was not considered a true *su(Hw)* mutant phenotype (Baxley et al., 2011; Harrison et al., 1993). To further examine the function of Su(Hw) in oogenesis, we performed a comprehensive analysis of the causes of female sterility in *su(Hw)* mutants including an evaluation of the aberrant nurse cell number phenotype. We used the *su(Hw)*^*e04061*^ mutant allele containing an insertion of a piggyBac transposon in the 5’ end of the second exon of *su(Hw)* (Baxley et al., 2011; Schoborg et al., 2013) as well as *su(Hw)*^*v*^, which carries a deletion of the *su(Hw)* promoter (Harrison et al., 1992). We observed an abnormal number of nurse cells in mutant egg chambers from both homozygous *su(Hw)*^*e04061*^ and trans-heterozygous *su(Hw)*^*v/e04061*^ mutants (Figure 1B-D). Therefore, we quantified the number of nurse cells per individual egg chamber in *su(Hw)*^*v/e04061*^ trans-heterozygous mutants using an antibody against lamin to stain the nuclear periphery of nurse cells. Mutant and wildtype stained ovaries were selected randomly, and egg chambers were analyzed using maximum projections of 20 Z-stack images that were collected with increments of 2 μm.

**Figure 1.**
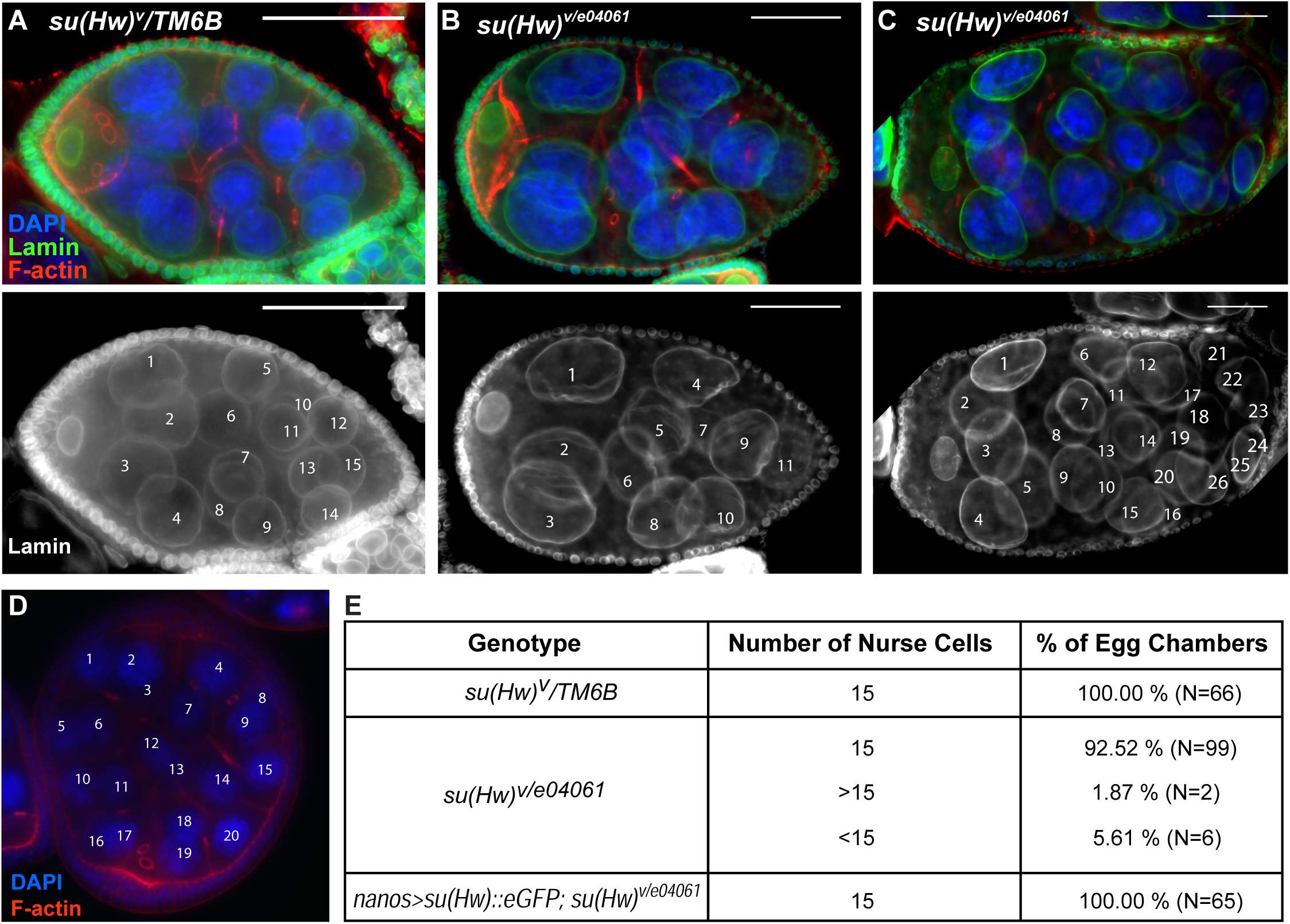
A fraction of egg chambers in *su(Hw)* mutants exhibit an irregular number of nurse cells. Wildtype and mutant egg chambers were stained with lamin antibody (green) and phalloidin (red), and nurse cell numbers were quantified. (A) A wildtype egg chamber with 15 nurse cells. (B) A *su(Hw)*^*v*/*e04061*^ egg chamber with less than 15 nurse cells. (C) A *su(Hw)*^*v*/*e04061*^ egg chamber with more than 15 nurse cells. (D) More than 15 nurse cells in *a su(Hw)*^*e04061*/*e04061*^ egg chamber. Scale bars are 50 μm. (E) *su(Hw)*^*v*^/*TM6B, su(Hw)*^*v*/*e04061*^, and *su(Hw)*::*eGFP, nanos*-*GAL4*; *su(Hw)*^*v*/*e04061*^ rescued individual egg chambers were documented in a table. The difference between wildtype and mutant as well as mutant and rescued flies are statistically significant (p<0.05, Fisher’s exact test). (F) The number of ring canals in *su(Hw)*^*v/e04061*^ mutants with greater than or less than 15 nurse cells were documented.

Results show that in addition to egg chambers with an increased number of nurse cells, some trans-heterozygous *su(Hw)*^*v/e04061*^ egg chambers also have a reduced number of nurse cells. Overall, 7.48% of the *su(Hw)*^*v/e04061*^ mutant population of egg chambers had an irregular number of nurse cells. Approximately 2% of egg chambers had more than fifteen nurse cells (Figure 1 E). This phenotype may arise from defective development of follicle cells, which may lead to fused chambers (Mata et al., 2000). Alternatively, this phenotype may result from additional mitotic divisions of cystoblasts occurring in the early germarium. Interestingly, 5.61% of egg chambers were found to have a reduced number of nurse cells, a phenotype that may indicate incomplete mitotic divisions in the germarium stage. This reduced nurse cell number phenotype upon loss of Su(Hw) was previously unreported and overall, the irregular nurse cell number in *su(Hw)* mutant ovaries compared with wildtype is statistically significant (p<0.05, Fisher’s exact test). Counting the number of ring canals connecting nurse cells with the oocyte provides an indication of whether egg chambers with an irregular nurse cell number originate from incomplete cell divisions, extra cell divisions, or by egg chamber fusions (Wang et al., 2010). After counting the number of ring canals contacting the oocyte in egg chambers with less than 15 nurse cells, we found that 33% contained less than 4 rings, suggesting incomplete nurse cell divisions. In 67% of egg chambers with less than 15 nurse cells, the number of rings was 4, suggesting cell death may have occurred in these chambers. Finally, supernumerary egg chambers always have more than 4 ring canals in their oocyte, suggesting these chambers had extra cell divisions (Figure S1).

To confirm that this phenotype is caused by the lack of Su(Hw) in the germline, we quantified nurse cell number in *su(Hw)*^*v/e04061*^ trans-heterozygous females expressing a Su(Hw):*eGFP* transgene in the germline driven by a *nanos*-*GAL4* driver (P{nos-*GAL4*::VP16}). The *nanos*-*GAL4* driver directs the expression of *GAL4* throughout all the stages of oogenesis (Rorth, 1998; Van Doren et al., 1998). We have previously shown that *su(Hw)::eGFP* completely rescues insulator activity in *su(Hw)* mutants (Schoborg et al., 2013) and partially rescues the fertility of these mutants when expressed in the germline (Hsu et al., 2015). Our results show that all 65 egg chambers analyzed in ovaries from females expressing the rescue construct contain the expected number of 15 nurse cells (Figure 1 E). Taken together, these observations suggest that loss of *su(Hw)* causes an irregular number of nurse cells.

### Microtubules are disorganized in *su(Hw)* mutant egg chambers

Abnormal nurse cell number in egg chambers occurs in loss of function mutants of genes encoding proteins involved in piRNA-related pathways, including *rihno* and *maelstrom* (*mael*) (Sato et al., 2011; Volpe et al., 2001). *mael* encodes a *γ*-tubulin associated protein involved in the proper positioning of the MTOC, which is required to determine oocyte polarity and the precise localization of specific mRNAs within the *Drosophila* oocyte (Clegg et al., 2001; Clegg et al., 1997; Sato et al., 2011). Microtubule organization is critical at various stages of oogenesis. In stage one, formation of the MTOC, a structure with concentrated *α*-tubulin at the posterior of the oocytes, is required for oocyte differentiation. In stages 3 through 6, a microtubule array is extended from the MTOC through ring canals to the neighboring nurse cells. This polarized network of microtubules is required for intercellular transport from nurse cells to the oocyte. During stage 7 the microtubule network is reorganized, causing a shift in the polarity of the MTOC from posterior to anterior, and the growing microtubule network positions the oocyte nucleus to the anterior corner (Steinhauer and Kalderon, 2006; Theurkauf et al., 1992).

To assess whether *su(Hw)* mutants also show microtubule disorganization in addition to an irregular number of nurse cells, we used an anti *α*-tubulin antibody that allows detection of microtubule networks in egg chambers (Theurkauf et al., 1992). We found that, under our experimental conditions, MTOCs form properly in wildtype ovaries, exhibiting the typical concentration of *α*-tubulin at the posterior of the oocyte in the germarium (Figure 2 C and G). However, in *su(Hw)* mutants, the *α*-tubulin signal is weaker, more diffuse, and less concentrated at the MTOC (Figure 2 A and E). This phenotype was specific to *su(Hw)* mutants, as we did not observe the same phenotype in the *mod(mdg4)*^*u1*^ mutant (Figure 2 B and F).

**Figure 2.**
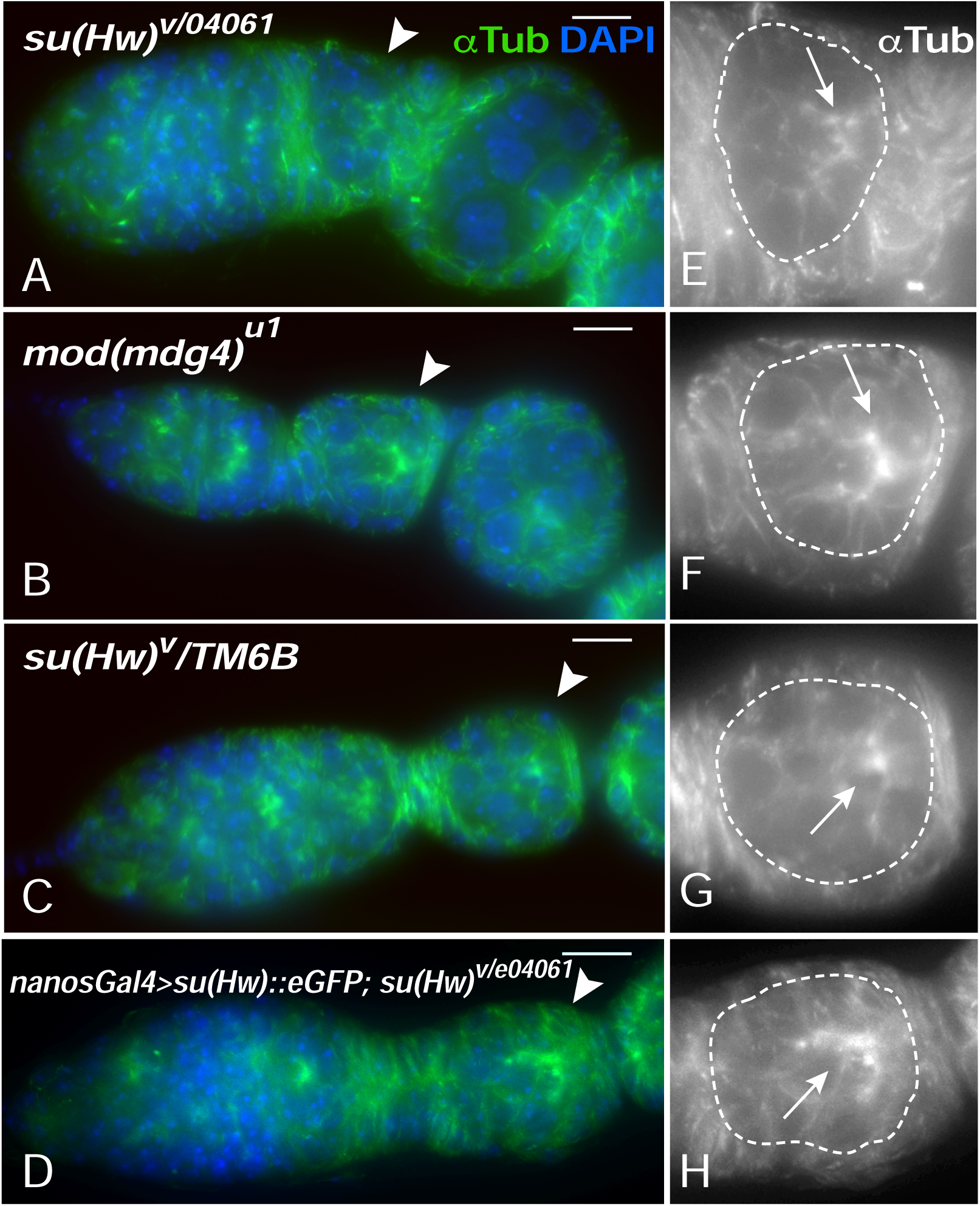
The MTOC is impaired in *su(Hw)* mutants. Microtubules are labeled with α-tubulin antibody (green). (A-D) Germarium stage egg chambers with different genotypes. Arrowheads indicate stage 1 egg chambers. (E-H) Egg chambers are magnified showing a dimmer and less bright MTOC (white arrowheads) in *su(Hw)*^*v*/*e04061*^ mutant chambers. Scale bars are 10μm.

Additionally, we were able to rescue the MTOC defect in the germarium of *su(Hw)* mutant ovarioles by ectopic expression of *su(Hw*)::*eGFP* driven by the *nanos*-*GAL4* driver (Figure 2 D and H). These data suggest that proper formation of the MTOC is impaired upon loss of Su(Hw) and imply that the microtubule network is disorganized and may not function efficiently to facilitate egg chamber development. For example, it is well established that defects in MTOC regulation that affect establishment of polarity in early oogenesis result in the disruption of Grk signaling in later stages of oogenesis (Khurana and Theurkauf, 2010).

### Gurken is mislocalized in the oocyte of *su(Hw)* mutant egg chambers

Embryo dorsoventral patterning is determined by the key axis-determining mRNAs of *gurken* (*grk*), *oskar* (*osk*), and *bicoid* (*bcd*), which are transported along microtubules to specific locations within the oocyte (Kugler and Lasko, 2009). To test whether defective microtubule organization in *su(Hw)* mutants could also affect axis determination, we used anti-Grk antibodies to detect the localization of Grk, a *Drosophila* transforming growth factor *α* (TGF *α*) protein, which is important for dorsoventral determination of embryos (Neuman-Silberberg and Schupbach, 1993). *grk* mRNA requires transportation to the oocyte prior to translation, and Grk protein specifically localizes at the posterior of the oocyte during stage 6 (Figure 3 A and B).

**Figure 3.**
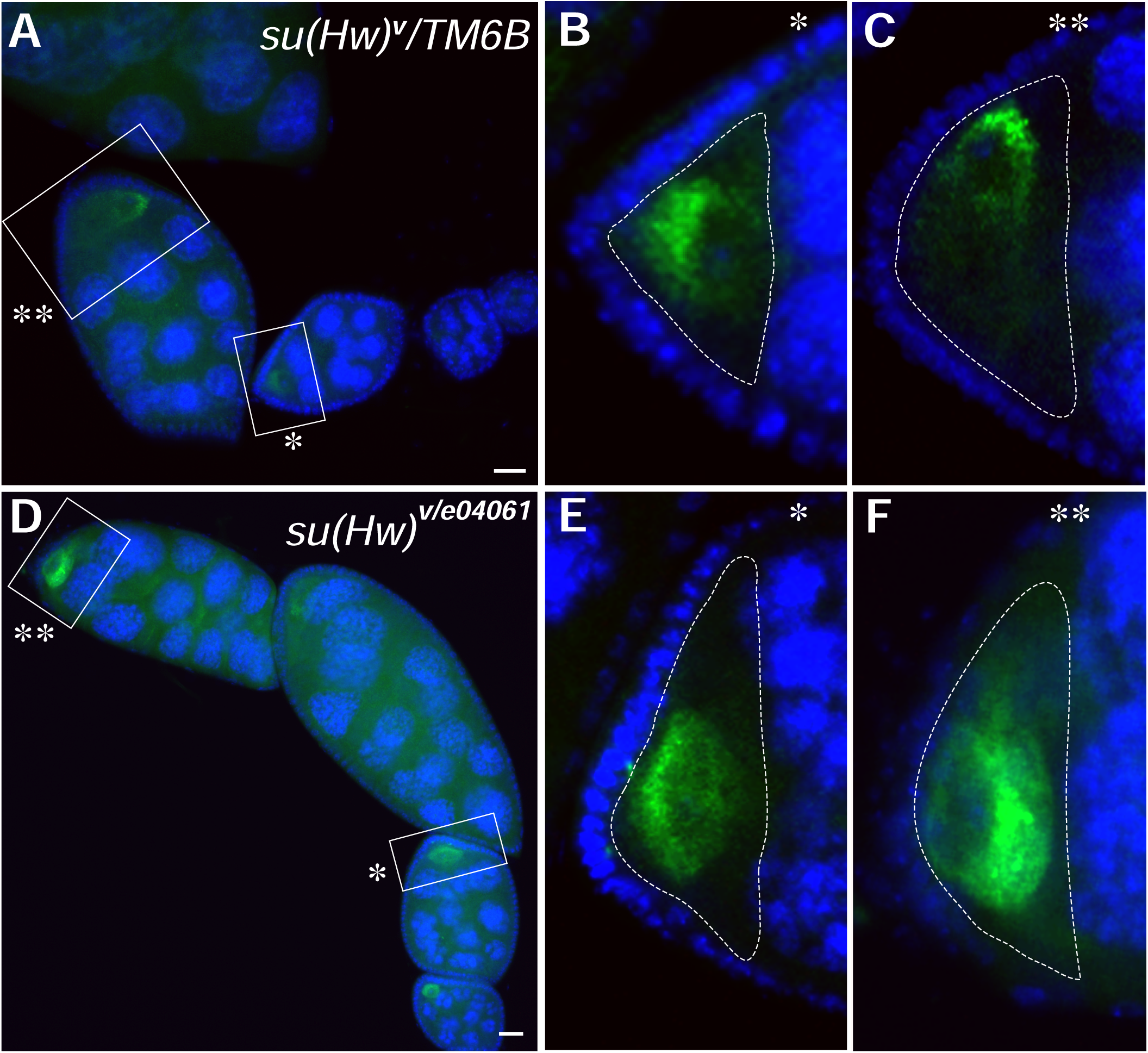
Gurken mislocalizes in *su(Hw)* mutant egg chambers. Grk was labeled in green and its location was monitored at stage 6 (*) and 9 (**) egg chambers. (A-C) In wildtype, Grk localizes at the posterior of the oocyte at stage 6 (inset B) and translocates to the dorsal-anterior corner at stage 9 (inset C). (D-F) Grk mislocalizes to the oocyte anterior at stage 9 in *su(Hw)*^*v*/*e04061*^ mutant. E and F are insets from D.

Later, as the oocyte relocates to the anterior-dorsal corner, Grk gradually moves to the corner and forms a crescent shape around the oocyte nucleus at stage 9 (Figure 3 C). Our data shows that in *su(Hw)* mutants, Grk fails to translocate to the anterior-dorsal corner of the oocyte in 95% (N=20) of the egg chambers at stage 9, compared with 100% correct localization in wildtype (N=16) (p<0.001, Fisher’s exact test) (Figure 6). Mislocalization of Grk normally leads to failure of the oocyte to signal follicle cells to determine dorsal fate, consequently interrupting the axis plan of the developing egg chamber. Our data suggest that loss of Su(Hw) causes defective formation of the microtubule network in developing egg chambers and the subsequent mislocalization of Grk.

### Non-meiotic DNA damage occurs during oogenesis in *su(Hw)* mutants

Disorganized microtubules and mislocalized Grk are oogenesis phenotypes observed in mutants of the *spindle* class genes (Ghabrial et al., 1998) and the piRNA pathway (Chen et al., 2007; Klattenhoff et al., 2007; Sato et al., 2011). In these mutants, cells lose the ability to repair DNA damage or to repress retrotransposon activity, generating an excess of DNA double-strand breaks (DSBs) such that unrepaired DSBs activate the ATR/Chk2 dependent DNA-damage signaling pathway in the female germline (Khurana and Theurkauf, 2010). Therefore, our results suggest that DNA damage signaling pathways are activated in *su(Hw)* mutants, resulting in disorganization of the MTOC and Grk mislocalization in the oocyte.

In order to further investigate whether the ATR-mediated DNA damage pathway is activated by DNA damage in *su(Hw)* mutants, we performed immunostaining in ovaries using antibodies against phosphorylated Histone 2A variant (*γ*H2Av), a marker for DNA damage. In *Drosophila* oocytes, DNA damage can be produced by three major sources: DNA damage induced by transposable element activity, DNA damage induced by replication stress, and meiotic DSB induced by Spo11, a nuclease required for meiotic recombination encoded by the *mei-w68* gene in *Drosophila*. These meiotic DSBs are normally repaired before developing egg chambers reach stage one (Jang et al., 2003; McKim and Hayashi-Hagihara, 1998; Mehrotra and McKim, 2006).

To characterize whether the loss of *su(Hw)* induces DNA damage and whether the DNA damage is associated with the meiotic recombination pathway, we used *γ*H2Av to compare the DNA damage in wildtype, *su(Hw)* mutants, *mei-w68* mutants, and *mei-w68*; *su(Hw)* double mutants. In wildtype germaria, we observed naturally occurring DNA damage in dividing cystocytes (which later become nurse cells and pro-oocytes) in region 2, where homologous recombination takes place (Figure 4 A and B). As expected, *mei-w68* (Spo11) homozygous mutants showed no DSBs in region 2 of the germarium, indicating a lack of Spo11 activity and the absence of other significant factors inducing DNA damage. Remarkably, we found an excess of non-meiotic DSBs in region 2 nurse cells from the germarium of *su(Hw)* and *mei-w68* double mutants, demonstrating that DNA damage in *su(Hw)* mutants is not produced by Spo11, and is therefore of non-meiotic origin (Figure 4 B and D). Additionally, we observed that in wildtype ovaries, *γ*H2Av signal is absent after stage 5, as described in an earlier study (Joyce et al., 2011). Interestingly, the *γ*H2Av signal remains present until later stages in *su(Hw)* mutant egg chambers (Figure 4 I and J). This result strongly indicates that loss of Su(Hw) leads to formation and accumulation of non-meiotic DNA damage in female germline cells.

**Figure 4.**
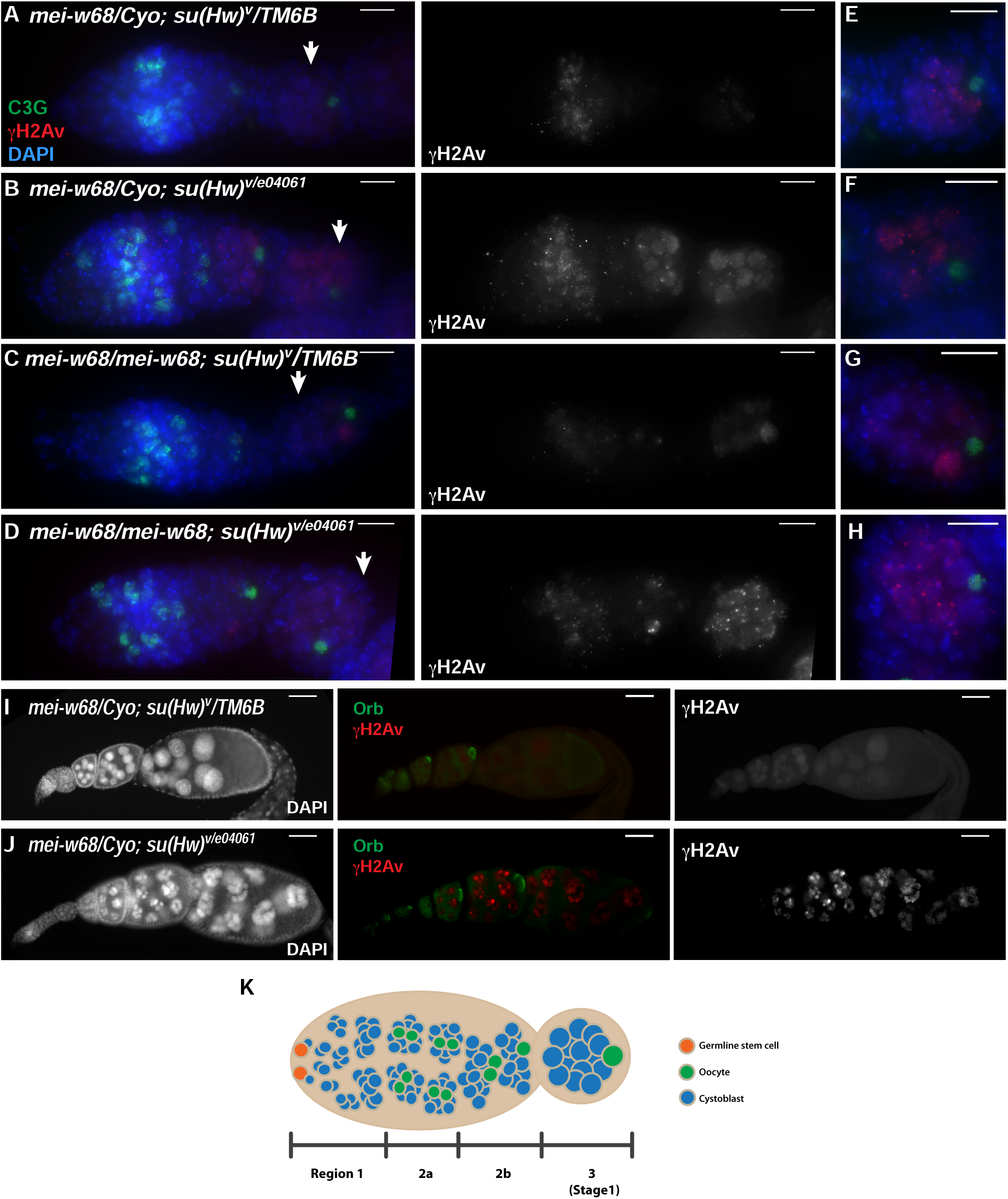
Non-meiotic DNA damage accumulates in *su(Hw)* mutant ovaries. DNA damage was visualized using an anti-*γ*H2Av antibody (red) in early germaria and later stage egg chambers. Anti-C(3)G (green), a synaptonemal complex protein, was used as a marker to identify pro-oocytes and regions 2a and 2b in ovarioles. Arrowheads point to stage 1 egg chambers. (A-H) Stage 1 egg chambers showing detail of nurse cells, oocytes, and *γ*H2Av foci in different genotypes. Scale bar in H is 10μm. (I-J) *γ*H2Av foci were detected in *su(Hw)*^*v*/*e04061*^ through oogenesis. The Orb staining in green indicates the location of the oocyte within the egg chambers and *γ*H2Av was observed to be relatively stronger in *su(Hw)*^*v*/*e04061*^ mutants compared to wildtype. The scale bar is 20 μm. (K) Diagram representing different stages of oogenesis in the *Drosophila* germarium.

### Su(Hw) does not play a major role in regulating global transposable element activity in the *Drosophila* germline

Su(Hw) strongly regulates transcription of the *gypsy* retrotransposon, and loss of the insulator activity reverses the phenotype of *gypsy* induced mutations at several loci in *Drosophila* (Harrison et al., 1993; Parkhurst and Corces, 1985; Parkhurst and Corces, 1986; Parkhurst et al., 1988). Additionally, global activation of transposable elements (TEs) in the germline triggers the activation DNA damage checkpoints most likely due to an excess of DNA damage, producing phenotypes similar to those described here (Chen et al., 2007; Khurana and Theurkauf, 2010; Klattenhoff et al., 2007; Mohn et al., 2014). Our data demonstrating the accumulation of non-meiotic DNA damage in female germline cells of *su(Hw)* mutants led us to ask whether TEs are over-expressed in the germline of these mutants.

We performed real-time RT-PCR to quantify the transcripts of TEs produced in wildtype and *su(Hw)*^*e04061*^ homozygous mutant ovaries. In both genotypes, egg chambers stage 9 and older were removed manually to eliminate differences in expression levels that could originate from differences in developmental stages between the two samples. Total RNA extracted from mutant and wildtype ovaries was used for real-time RT-PCR. We used 17 primer sets specifically designed to determine the expression of 17 different TEs. Additionally, we used primers for *rp49* as an internal control. Our selection of TEs includes germline and somatic specific transposons and covers TEs with long-terminal repeats (LTR) as well as non-LTR elements.

The expression levels of 17 TEs in mutant ovaries were compared to wildtype in a fold-change graph (Figure 5). Consistent with previous studies, our data show that the transcription levels of *gypsy* as well as *stalker4* are reduced significantly in *su(Hw)* mutants. Compared with the transcription levels of TEs in spindle mutants, which can reach levels of more than 100-fold increase, and considering that the majority of TEs tested showed no significant change in transcription levels, it seems unlikely that TEs are the cause of significant DNA damage in *su(Hw)* mutants.

**Figure 5.**
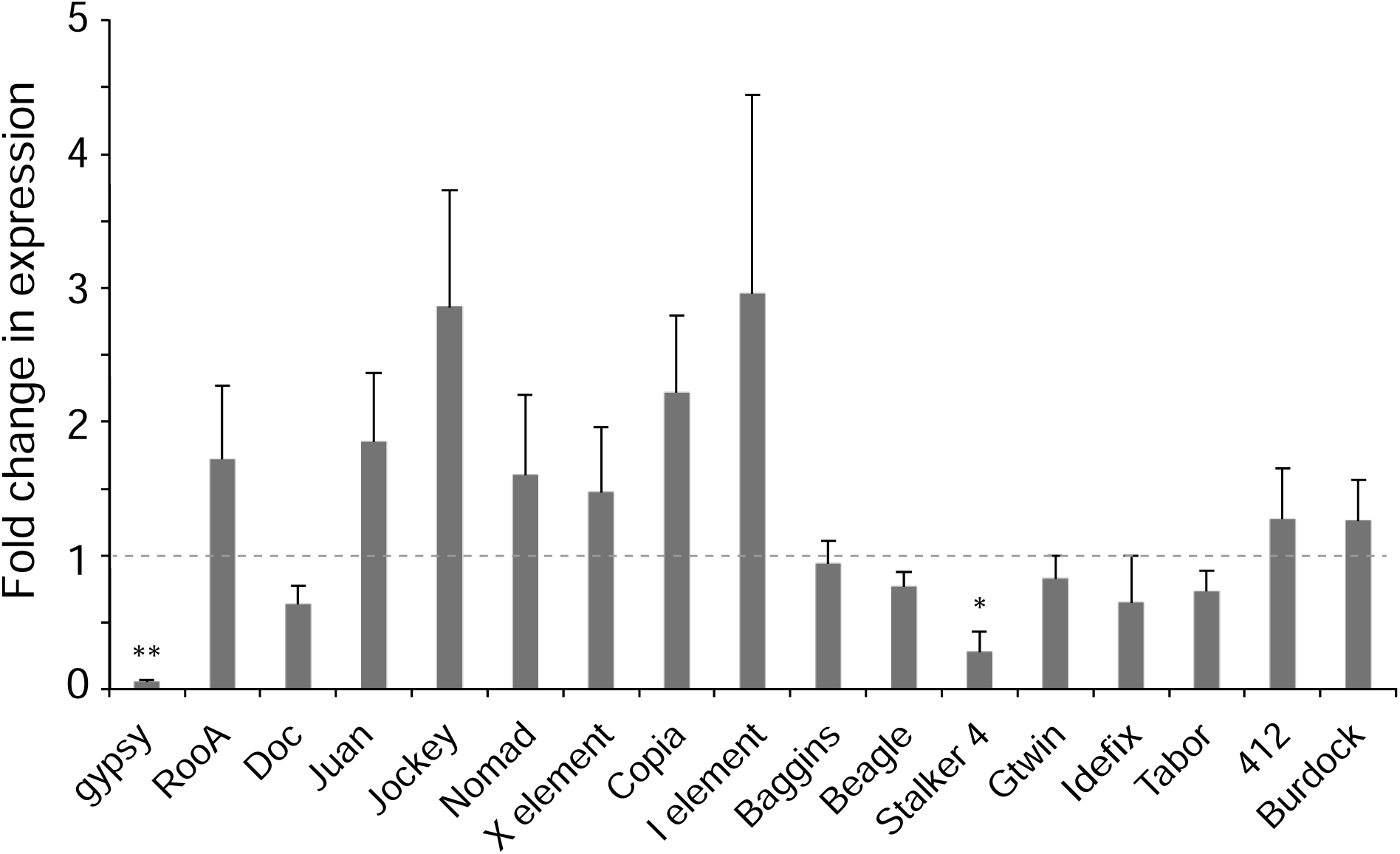
Expression of transposable elements does not increase significantly in *su(Hw)* mutant ovaries: Transcript levels of seventeen transposable elements were quantified by real-time PCR and were normalized using rp49 as a control. Fold change values represent the relative expression of mRNA in ovaries from *su(Hw)*^*e04061*^ homozygotes compared with ovaries from *su(Hw)*^*e04061*^/*TM6B* heterozygotes. One asterisk indicates P < 0.05 and two for P<0.0001.

### Loss of Su(Hw) activates a DNA damage checkpoint during oogenesis

Because of the similarity between spindle class and piRNA mutants with *su(Hw)* mutant phenotypes and the elevated levels of *γ*H2Av seen in *su(Hw)* mutant egg chambers, we hypothesize that *su(Hw)* mutations activate a DNA damage checkpoint during oogenesis. We first tested whether *su(Hw)* mutants activate a DNA damage signaling pathway by asking whether oocyte development is restored in females double mutant for *su(Hw)*^*v/e04061*^ and the *Drosophila* ATR allele, *mei-41*^*D5*^ (Brodsky et al., 2004; Laurencon et al., 2003). Results show that although double mutant females remained sterile, 54% of egg chambers recovered correct positioning of Grk around the oocyte nucleus at stage nine, (N=24, p<0.001, Fisher exact test), and had proper enlargement of the developing oocyte in stage nine and ten (Figure 6 C, H, and I). These results show that loss of ATR function partially recovers oocyte development in *su(Hw)*^*v/e04061*^ mutants and suggests that loss of Su(Hw) function triggers a DNA-damage response through an ATR-dependent pathway.

**Figure 6.**
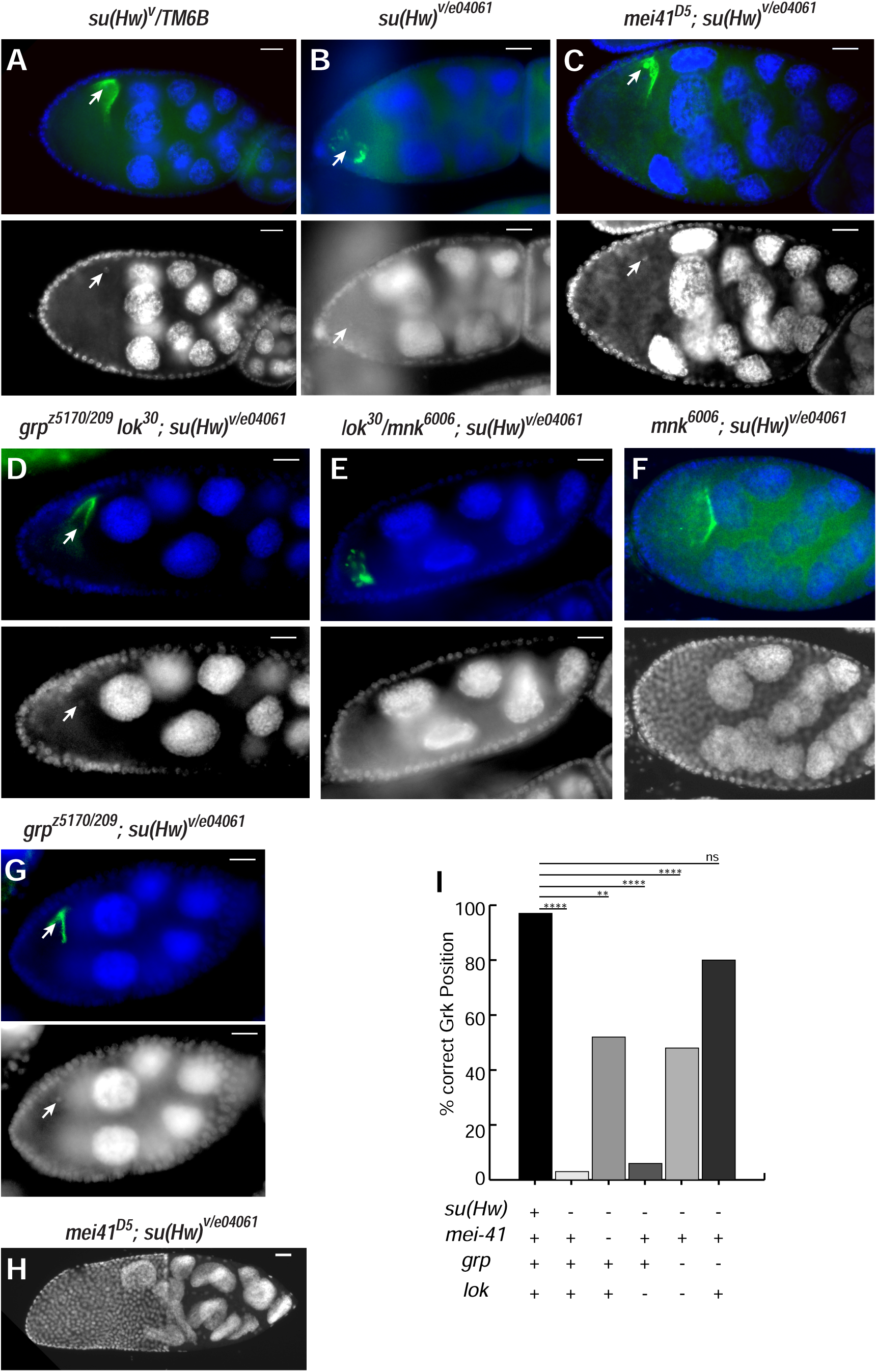
Oogenesis failure in *su(Hw)*^*v*/*e04061*^ mutants is partially rescued by *mei-41*^*D5*^ (ATR) and *grp*^*z5170*/*209*^ (Chk1) mutations. Grk (green) was detected in egg chambers. (A-G) A *su(Hw)*^*v*^/*TM6B* egg chamber with normal Grk signaling (A), *su(Hw)*^*v*/*e04061*^ egg chamber with Grk mislocalization (B), *mei41*^*D5*^; *su(Hw)*^*v*/*e04061*^egg chamber with rescued Grk localization (C), *grp*^*z5170*/*209*^*lok*^*30*^;*su(Hw)*^*v*/*e04061*^ egg chamber with rescued Grk localization (D), *lok*^*30*^*mnk*^*6006*^; *su(Hw)*^*v*/*e04061*^egg chamber with Grk mislocalization (E), a *mnk*^*6006*^; *su(Hw)*^*v*/*e04061*^ egg chamber with Grk mislocalization (F), and a *grp*^*z5170*/*209*^;*su(Hw)*^*v*/*e04061*^ egg chamber with rescued Grk localization (G) are shown. (H) A *mei-41*^*D5*^, *su(Hw)*^*v*/*e04061*^ double mutant egg chamber at stage 10. Scale bars are 20μm. (I) Graph shows percentage of correct Grk localization in egg chambers from each genotype.

Chk1 (Grp) and Chk2 (Mnk, Lok), two highly conserved downstream kinases in ATR/ATM DNA damage signaling, are both phosphorylated in response to DNA damage and participate in cell cycle checkpoint activation (Cimprich and Cortez, 2008). However, mutations of *chk2*, but not *chk1*, recover the dorsoventral patterning defects associated with mutants of spindle and piRNA pathway genes (Chen et al., 2007; Ghabrial et al., 1998; Klattenhoff et al., 2007). To determine the role of DNA repair pathways in *su(Hw)* mutant ovaries, we first tested the ability of *chk1* (*grp*^*z5170/209*^) *chk2* (*lok*^*30*^) double mutant flies to rescue the mutant phenotypes of *su(Hw)*^*v/e04061*^ ovaries. We found that *grp* ^*z5170/209*^ *lok*^*30*^ double mutants were able to rescue defective Grk localization, but were unable to rescue the lack of fertility in *su(Hw)* mutants (Figure 6 D). To test whether the loss of Su(Hw) activates a DNA damage response specifically mediated by ATR/Chk2, we generated double mutant flies using *su(Hw)*^*v/e04061*^ and *chk2* (*lok*^*30*^, *mnk*^*6006*^) mutations (Brodsky et al., 2004) and tested whether a mutation in *chk2* is able to rescue Grk localization in *su(Hw)* mutants. The results showed that neither fertility nor Grk localization were rescued in these double mutants (Figure 6 E and F). Finally, we generated double mutant flies using *su(Hw)*^*v/e04061*^ and *chk1* (*grp*^*z5170/209*^) mutations. *grp*^*z5170/209*^; *su(Hw)*^*v/e04061*^ double mutants were able to rescue the Grk localization phenotype in 80% of egg chambers analyzed, but were unable to rescue the sterility phenotype (Figure 6 G and I). These findings show that the spindle-like phenotypes caused by loss of Su(Hw) are dependent on Chk1 (*grapes*) mediated checkpoint activity, downstream of the ATR mediated DNA-damage pathway, and are independent of Chk2 (*mnk, lok*). Since ATR/Chk1 activate a checkpoint in response to replication stress (Blythe and Wieschaus, 2015; Fogarty et al., 1997; Sibon et al., 1997), our results suggest that mutations of *su(Hw)* cause replication stress in developing egg chambers.

### Misexpression of Su(Hw) in mutant ovaries causes dorsoventral defects in eggshells

Our results have revealed that Su(Hw) may have a role in maintaining genome integrity by preventing replication stress in nurse cells. Accordingly, loss of Su(Hw) causes a significant increase in DNA damage, followed by defects in microtubule organization, egg chamber formation, and mislocalization of Grk. These phenotypes can be rescued by *mei-41*^*D5*^ and *grp*^*z5170/209*^, mutant alleles of ATR and Chk1 respectively, suggesting that the defects originate from the activation of checkpoints in response to DNA damage.

In addition to these phenotypes, homozygous mutant females for spindle class genes produce embryos with dorsoventral defects in their eggshells which result from mislocalization of Grk and other morphogens in the developing oocyte (Ghabrial et al., 1998; Ghabrial and Schupbach, 1999). Egg chamber development in *su(Hw)* mutants is arrested at mid-oogenesis, preventing the opportunity to test eggshells for the formation of dorsoventral defects. Interestingly, we have shown that *su(Hw)*::*eGFP* expression driven by *nanos*-*GAL4* (P{nanos-*GAL4*::VP16}) not only can rescue MTOC in egg chambers, (Figure 2) but also partially restores fertility in mutant flies. These rescued females laid a reduced number of eggs (∼41% of normal females) (Hsu et al., 2015), and only less than 25% of these embryos produced viable adults, indicating that the spatio-temporal expression of *nanos*-*GAL4*; *su(Hw)*::*eGFP* is not sufficient to fully restore Su(Hw) activity in all egg chambers from mutant ovaries.

Eggshells produced by *nanos*-*GAL4*; *su(Hw)*::*eGFP* partially rescued females provide an opportunity to test whether loss of Su(Hw) also induces the formation of dorsoventral defects. We performed a quantitative analysis of the eggshell morphology produced by *nanos*-*GAL4*; *su(Hw)*::*eGFP* females. Results show that all embryos that failed to develop displayed eggshells with dorsoventral axis defects (Figure 7A). We categorized these phenotypes as previously described in (Ghabrial et al., 1998): Type I: eggshells with two wildtype appendages, Type II: eggshells showing two short appendages, Type III: eggshells showing a single appendage, and Type IV: eggshells with no appendages. The number of each type of eggshell is shown in Figure 7B. Type I eggshells are only found in viable embryos and have no noticeable developmental defects through adult stage. These results show that in a large number of egg chambers expressing *su(Hw)*::*eGFP* driven by *nanos*-*GAL4*, oogenesis defects are only partially rescued, and suggest that these chambers activate a spindle-like checkpoint that leads to dorsoventral transformations.

**Figure 7.**
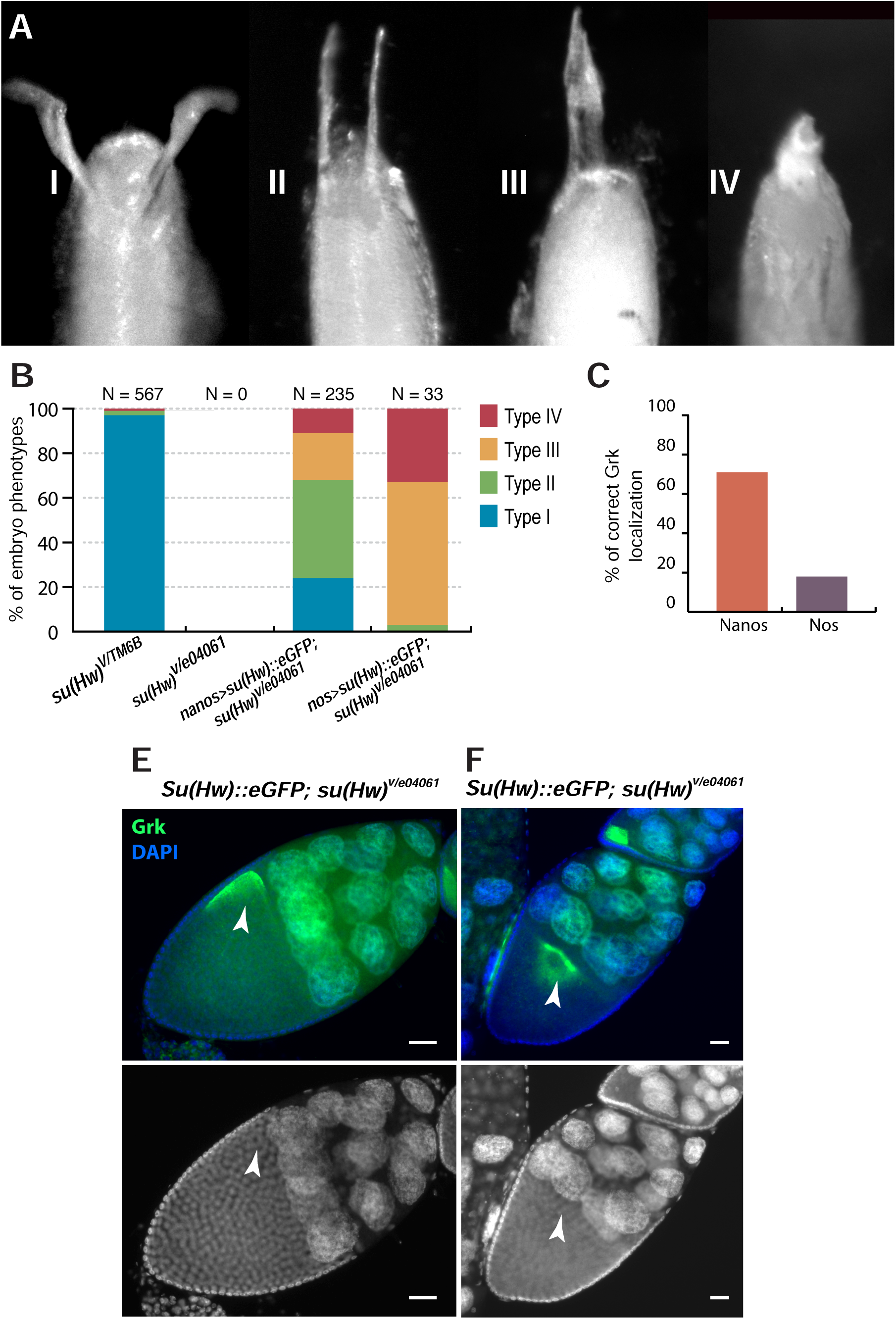
Expression of Su(Hw) in germline cells rescues female fertility and partially rescues embryo development. Wildtype virgin females and rescued females expressing either *nanos*-*GAL4* driven *su(Hw)*::*eGFP* or *nos*-*GAL4* driven *su(Hw)*::*eGFP* were crossed with wildtype male flies for three days and eggs were collected. (A) Different dorsoventral phenotypes of embryos in the offspring were categorized as Type I to IV, and the percentage of each category of embryos is shown in the stacked column graph. (B) *su(Hw)*^*v*^/*TM6B* female flies produced 97% type I, 2% type II, and 1% type IV embryos; however *nanos*-*GAL4* driven *su(Hw)*::*eGFP* rescued flies have only 24% type I (two normal appendages), 44% type II (two short appendages), 21% type III (one appendage), and 11% type IV (no appendage), while *nos*-*GAL4* driven *su(Hw)*::*eGFP* rescued flies have 0% type I, 3% type II, 64% type III, and 33% type IV. Sterile *su(Hw)* mutant females were used as negative control. (C) Grk localization in rescued egg chambers. In *nanos*-*GAL4* driven *su(Hw)*::*eGFP* rescued flies (N=7), 71 % of egg chambers had correct Grk localization, but in *nos*-*GAL4* driven *su(Hw)*::*eGFP* rescued flies only 18% egg chambers had normal Grk localization (N=11). (D and E). Grk localization in stage 9 egg chambers from rescued females. Arrowheads point to Grk. Green signal in nurse cell nuclei, in D, corresponds to GFP signal from *su(Hw)*::*eGFP*.

Surprisingly, it has been recently shown that heterozygous mutations in the *Rbp9* gene (*Rbp9*^*P2775/+*^, *su(Hw)*^*2/v*^) are able to rescue the sterility of *su(Hw)* mutant females (Soshnev et al., 2013). However, embryos produced by these females show dorsoventral transformations in their eggshells. Soshnev and collaborators conclude that transformations are produced because of the lack of Su(Hw) expression in follicle cells, suggesting that expression of Su(Hw) in follicle cells is necessary to allow establishment of communication between Gurken and follicle cell receptors. In experiments described here, we used a *nanos*-*GAL4* driver to express Su(Hw) in nurse cells. This driver induces transgene expression exclusively in the germline, showing expression peaks at the germarium and at stage 9, but does not activate expression in follicle cells (Rorth, 1998; Van Doren et al., 1998). We tested expression of Su(Hw)::*eGFP* in follicle cells using antibodies against GFP, showing no detectable expression, yet 20% of the embryos were able to develop into normal adult flies, suggesting that expression of *su(Hw)* is not required in follicle cells to support correct dorsoventral patterning during oogenesis (Hsu et al., 2015). Additionally, we found that the source of ventralized phenotypes in *nanos-GAL4*; *su(Hw)*::*eGFP* eggshells can be attributed to mislocalization of Grk, since 29% of egg chambers from ovaries of these females also show mislocalization of Grk (Figure 7C and D).

We also generated *nos-GAL4*, Su(Hw)::*eGFP* flies, in which the driver is P{*GAL4*-nos.NGT}40), that highly express Su(Hw)::*eGFP* through oogenesis. In these flies, the sterility phenotype of *su(Hw)* mutant females was poorly rescued, but a large number of egg chambers achieved normal oocyte enlargement and were able to reach the latest stages of oogenesis. *nos-GAL4*, Su(Hw)::*eGFP*; *su(Hw)*^*v/e04061*^ flies laid 6.1% of expected eggs, compared to wildtype (Hsu et al., 2015). *Nanos*-*GAL4*::VP16 includes the *nos* 5’ and 3’ UTRs translation regulation sequences and appears to be a stronger driver than *GAL4*-*nos*.NGT, which includes a 3′UTR region from a tubulin mRNA (Tracey et al., 2000). Embryos from *nos-GAL4*, Su(Hw)::*eGFP* females never developed into larvae and all eggshells showed ventralization phenotypes with the more severe classes more frequently represented than in embryos from *nanos*-*GAL4*::VP16 driver (Figure 7B). Importantly, the severity of the ventralization phenotypes correlates with a higher frequency of mislocalization of Grk (82%) (Figure 7C and E), suggesting that defects stem from misexpression of Su(Hw) in oocytes and not from the lack of *su(Hw)* expression in follicle cells. Altogether, this data further supports our findings that expression of *su(Hw)* is required for genome stability during oogenesis and that loss of Su(Hw) results in the generation of DNA damage, followed by the activation of cell cycle checkpoints, mislocalization of Gurken, and dorsoventral patterning defects in eggshells.

### Monomethylated H4K20 and γH2Av accumulate in *su(Hw)* mutant ovaries

We have shown that loss of Su(Hw) causes an accumulation of DNA damage in nurse cells, but its origin remains uncertain since the damage is not derived from meiotic recombination and a substantial activation of TEs was not observed in these mutants. Given that, in addition to Spo11 and TE activity, a major contributing source of intrinsically produced DNA damage under normal conditions is replication stress (Zeman and Cimprich, 2014) and, because the spindle-like phenotype we observed is ATR/Chk1 dependent, we used monomethylated Histone 4 at lysine 20 (H4K20me1) as a marker to further test whether DNA damage in ovaries from *su(Hw)* female mutants is related to irregularities in replication. H4K20me1 has an important role in maintaining genome stability (Beck et al., 2012b; Jorgensen et al., 2013; Wu and Rice, 2011). Monomethylation of H4K20 is mediated by the PR-Set7/SET8 methyltransferase, and removal of PR-Set7 results in DNA damage and S phase arrest (Jorgensen et al., 2007). Conversely, constant expression of PR-Set7 causes accumulation of H4K20me1 at replication origins and results in replication stress due to re-replication (Jorgensen et al., 2007; Tardat et al., 2010b). In *Drosophila* S2 cells, depletion of PR-Set7 affects chromosome compaction and higher-order chromatin organization, also triggering the DNA damage response (Sakaguchi et al., 2012; Sakaguchi and Steward, 2007). Additionally, highly proliferating tissues, such as wing discs and salivary glands, are smaller in size and contain fewer cells in *pr-set7* mutants, due to improper cell division during development (Karachentsev et al., 2007; Karachentsev et al., 2005). Together, this evidence suggests that misregulation of H4K20me1 can alter chromatin organization and lead to genome instability in a manner dependent on the cell cycle and DNA replication.

We performed western blotting in ovaries using both anti-*γ*H2Av and anti-H4K20me1 antibodies. Our results show that, as expected, the amount of *γ*H2Av and the amount of H4K20me1 increased in *su(Hw)* mutant ovaries compared to wildtype (Figure 8 B). The increase in *γ*H2Av levels revealed by western blots confirms our earlier observation that high levels of non-meiotic DNA damage occur in *su(Hw)* mutant ovaries (Figure 8 A). We also showed that an additional mutation in ATR (*mei-41*^*D5*^), which partially rescues oogenesis in *su(Hw)*^*v/e04061*^, does not significantly reduce the levels of *γ*H2Av in mutant ovaries (Figure 8 A), suggesting that H2Av is likely phosphorylated also by ATM in *su(Hw)* mutant ovaries. In addition, the elevated amounts of H4K20me1 in mutant ovaries support the notion that DNA damage may originate from replication stress, which we propose is caused by the loss of Su(Hw). Altogether, our results suggest that DNA damage in *su(Hw)* mutants is caused by replication stress, which disrupts genome stability and eventually leads to checkpoint activation and contributes to the oogenesis phenotype of these mutants.

**Figure 8.**
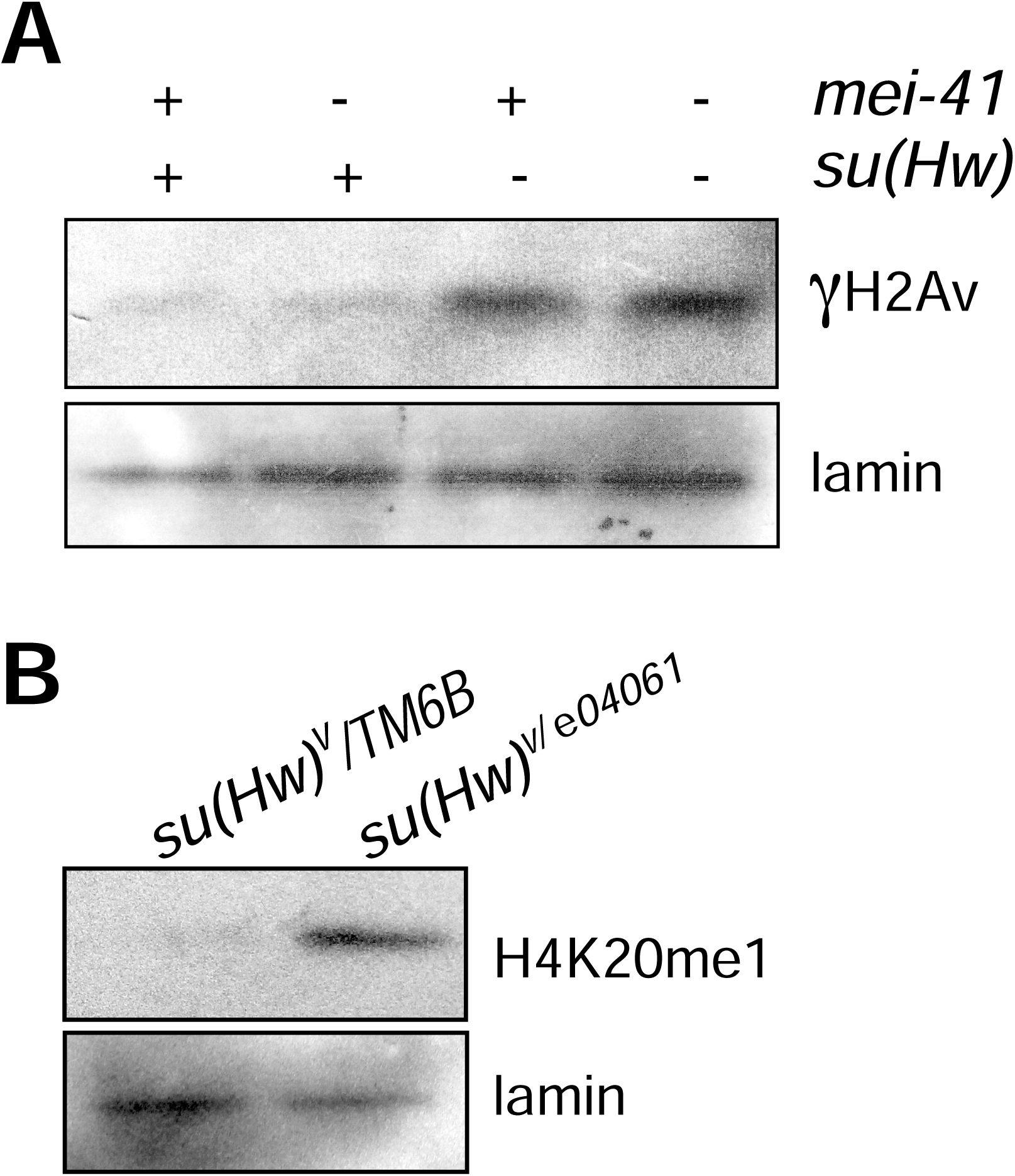
Levels of γH2Av and H4K20me1 are increased in *su(Hw)* mutant ovaries. (A-B) Western blot analysis of γH2Av and H4K20me1 in wildtype and *mei-41*^*D5*^ *su(Hw)* mutant ovary extracts. (A) Increased γH2Av is detected in *su(Hw)*^*v*/*e04061*^ and *mei-41*^*D5*^; *su(Hw)*^*v*/*e04061*^ double mutants, compared to wildtype (*su(Hw)*^*v*^/*TM6B*). Lamin was used as a loading control. (B) H4K20me1 levels increase in *su(Hw)*^*v*/*e04061*^ compared to wildtype (*su(Hw)*^*v*^/*TM6B*).

### Mutation of *su(Hw)* is associated with chromosomal aberrations in somatic tissues

Results showing spindle like phenotypes in *su(Hw)* mutant ovaries, and a partial rescue of this phenotype by mutations in *mei-41* and *grapes*, suggest Su(Hw) may play a role in the maintenance of genome stability and that lack of Su(Hw) function may lead to replication stress. However, these data did not show whether Su(Hw)’s role in genome maintenance is general or if this role is specific to the female germline. To address this question, we obtained metaphase spreads from third instar larval neuroblasts mutant for *su(Hw)*, and chromosomes were scored for the presence of chromosomal aberrations (CABs). Dividing neuroblasts provide an ideal opportunity to score a large number of cells and have previously been used to quantify the occurrence of CABs (Gatti and Goldberg, 1991). Screening larval brains from homozygous *su(Hw)*^*e04061*^ larvae reveals neuroblasts with either normal karyotypes (Figure 9, A1) or karyotypes showing CABs (Figure 9, A2-6). Different types of aberrations were noted in this genetic background, including many with chromatid or isochromatid deletions. Numerous cells of this genotype carried more than one aberration (Figure 9, A4), with some genomes appearing largely fragmented (Figure 9, A5-6). The frequency of CAB occurrence per brain between genotypes was quantified over multiple brains, revealing a significant increase in the average ratio of cells per brain that contain CABs in the homozygous *su(Hw)*^*e04061*^ background (11.9%, Figure 9 B and C) as compared to the wildtype strain Oregon R (OR) (p=0.004). The frequency of CABs observed in OR is 0.7%, close to previously reported values for wildtype CAB frequency (Mengoli et al., 2014; Merigliano et al., 2017), and is close in value to that obtained from the *su(Hw)*^*e04061*^/TM6B heterozygote. Examination of brains from a trans-heterozygous *su(Hw)*^*v/e04061*^ genotype showed a similar increase in CAB frequency over OR as the *su(Hw)*^*e04061*^ homozygote (p=0.0298), implying that this phenotype is not due to second-site mutations in the *su(Hw)*^*e04061*^ genetic background. This suggests that *su(Hw)* has a role in maintaining chromosome integrity and that half the wildtype dosage of *su(Hw)* is sufficient to prevent chromosomal aberrations from occurring.

**Figure 9.**
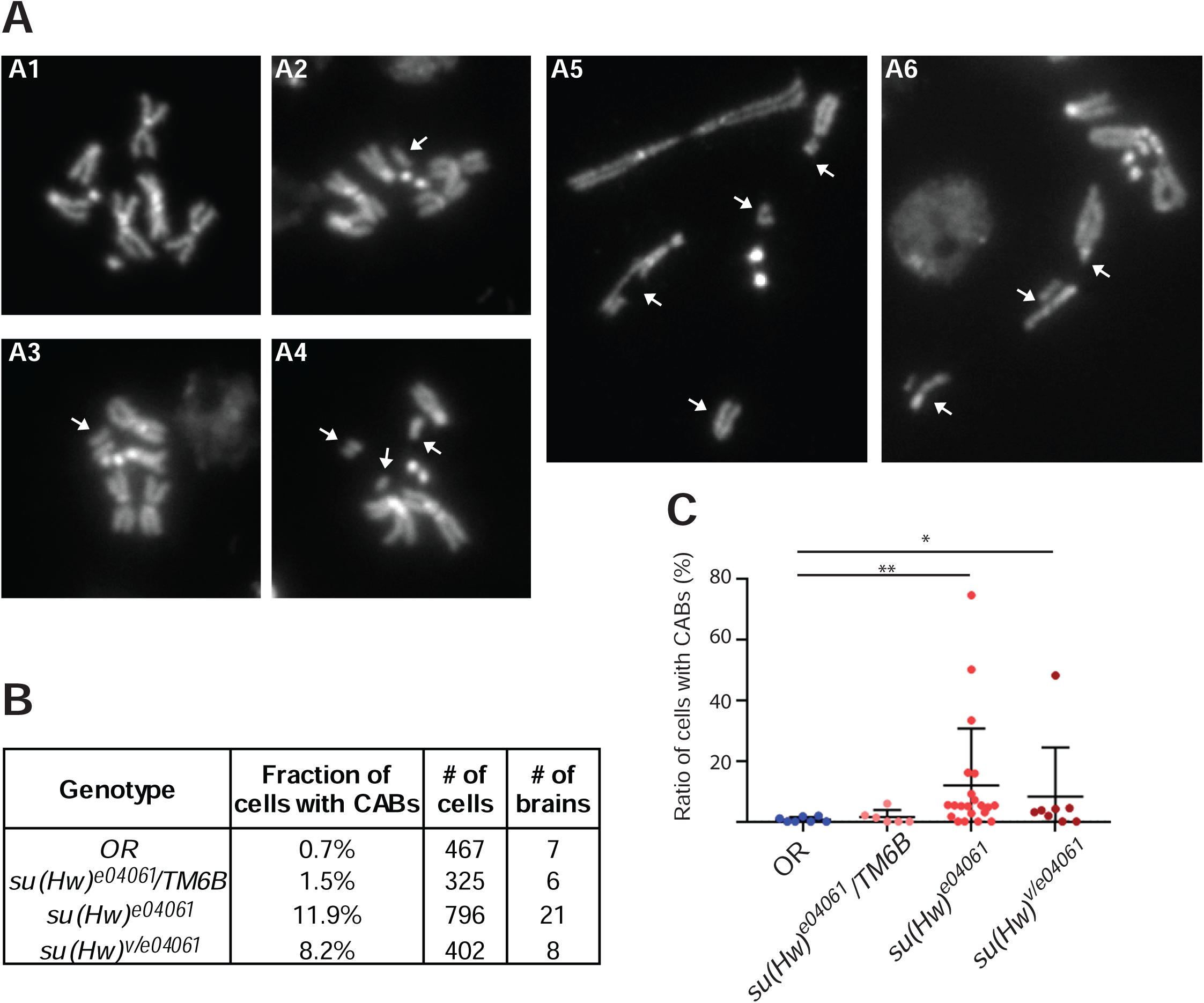
Mutation of *su(Hw)* is associated with chromosomal aberrations (CABs). (A) Examples of wild type chromosomes (A1, labelled with chromosome numbers) and those with chromosomal aberrations (A2-A6). Chromatid deletions (A2) and isochromatid deletions (A3) are frequently found in *su(Hw)* mutant backgrounds. (A4-A6) Genomes showing multiple CABs per cell. White arrows denote the sites of chromosomal breaks. (B) The fraction of cells showing CABs from each brain according to genotype. (C) A significant frequency of chromosomal aberrations is seen in *su(Hw)*^*e04061*^ homozygotes and *su(Hw)*^*v/e04061*^ trans-heterozygotes but not for wild-type or *su(Hw)* ^*e04061*^ heterozygotes. Error bars represent one standard error. * p < 0.05, ** p < 0.005.

## Discussion

This study has uncovered a novel role for Su(Hw) in the maintenance of genome stability. This conclusion is supported by data showing that a DNA damage response is activated in *su(Hw)* mutant egg chambers as well as by the occurrence of chromosomal aberrations in dividing neuroblasts from *su(Hw)* mutant larval brains. Traditionally, gene mutations that lead to the activation of DNA damage response pathways in the germline of *Drosophila* females are recognized by the formation of dorsoventral patterning defects in eggshells (Abdu *et al*., 2002; Ghabrial *et al*., 1998). In *su(Hw)* mutant ovaries, however, oogenesis arrests and oocytes do not fully develop, preventing direct observation of whether dorsoventral patterning defects in eggshells are an element of the phenotype. This, in combination with the circumstance that all other phenotypes associated with the production of elevated intrinsic DNA damage cannot be directly observed without the appropriate experimental analysis, may explain why the phenotype of *su(Hw)* mutations has rarely been previously associated with DNA damage or the DNA damage response (Lankenau et al., 2000).

Here, we have shown that an irregular number of nurse cells, MTOC disorganization, and Grk mislocalization are defects in *su(Hw)* mutant egg chambers. We also show that germline cells of *su(Hw)* mutants undergo an excessive accumulation of H2Av phosphorylation. Furthermore, our data suggests that DNA damage activates an ATR mediated DNA response signaling pathway, which may activate checkpoints leading to a failure of MTOC formation and Grk mislocalization. The occurrence of these phenotypes also provides an explanation for our observation that females ectopically expressing Su(Hw) systematically produce eggshells with dorsoventral defects. These phenotypes are usually observed in mutants encoding DNA repair or piRNA pathway components (Figure 10) (Abdu et al., 2002; Chen et al., 2007; Ghabrial and Schupbach, 1999; Gonzalez-Reyes et al., 1997; Klattenhoff et al., 2007; Staeva-Vieira et al., 2003).

**Figure 10.**
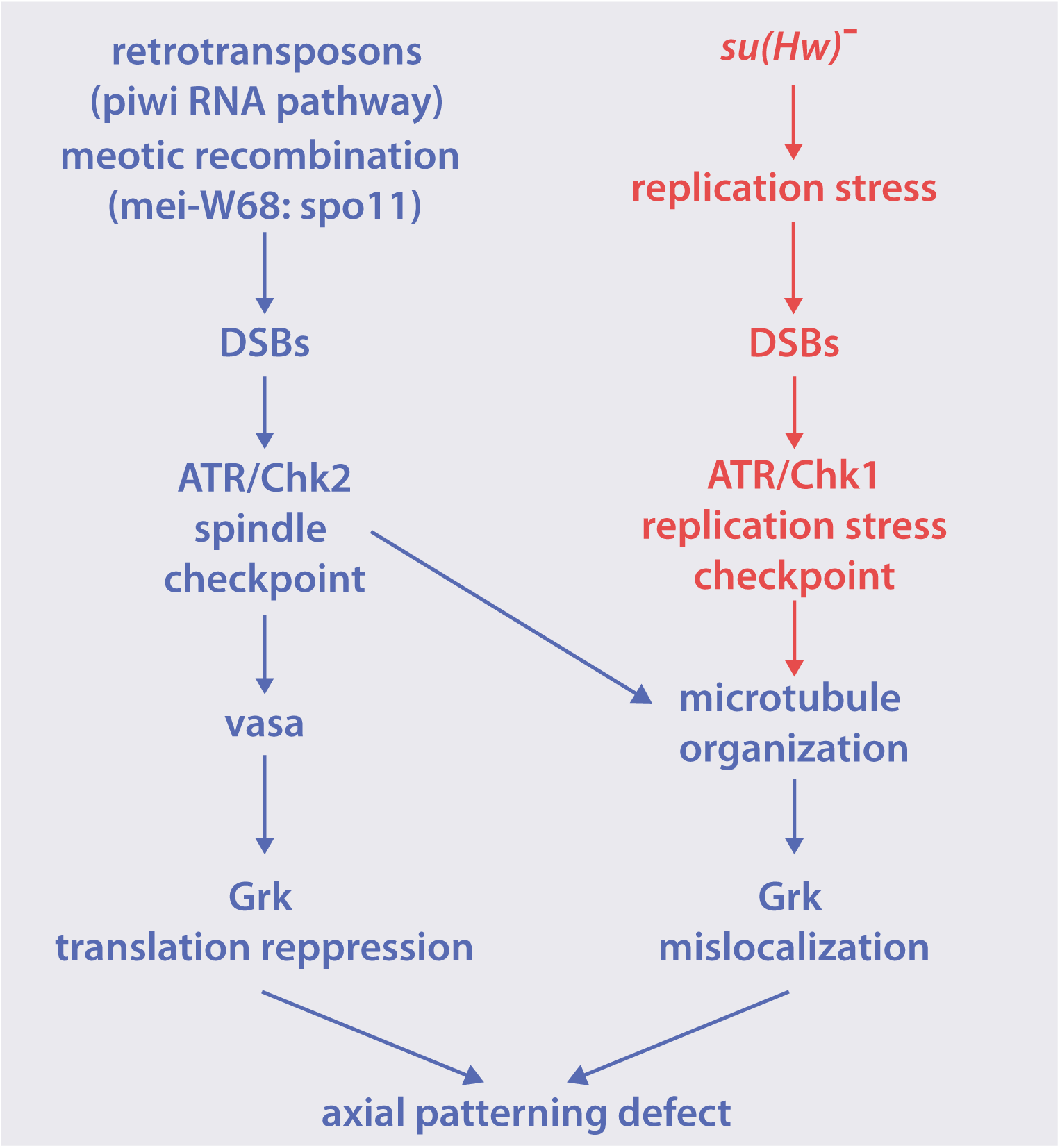
An ATR/Chk1 checkpoint activation pathway is activated during replication stress in ovaries from *su(Hw)* mutant females. Three known causes of DNA damage during oogenesis are shown: meiotic recombination, transposable elements, and replication stress. Mutation of genes encoding DSB repair proteins or helicase (spindle mutants, left) disable the mechanisms that repair meiotic DSBs generated at an early stage of oogenesis. Overactivated Transposable Elements (TEs) may produce an excess of DSBs. This type of DSBs is usually observed in piRNA pathway mutants. In both spindle mutants and piRNA pathway mutants, unrepaired DSBs activate an ATR/Chk2 dependent DNA damage checkpoint that leads to a sequence of developmental defects, including microtubule disorganization, Grk mislocalization, Grk repression, and patterning defects in eggshells. In *su(Hw)* mutant females, replication stress induces DNA damage during endoreplication (red pathway) and activates an ATR/Chk1 checkpoint that leads to Grk mislocalization and dorsoventral transformations.

Ataxia Telangiectasia-Related (ATR) and Ataxia Telangiectasia-mutated (ATM) are two highly conserved kinases with central roles in DNA repair, cell-cycle checkpoint progression, and cell fate determination (Sancar et al., 2004). In the DNA damage response, ATM primarily acts in repair of DSBs, whereas ATR is activated in response to various DNA lesions, particularly those generated by replication stress (Zeman and Cimprich, 2014). Both kinases are able to phosphorylate unique downstream effectors as well as a number of common targets, such as the human H2AX or the *Drosophila* equivalent H2Av. Phosphorylated H2AX (*γ*H2AX) in humans, or phosphorylated H2Av (*γ*H2Av) in *Drosophila*, is an indicator for ATR/ATM activity, and crosstalk between these two pathways occurs in response to DNA damage (Cimprich and Cortez, 2008; Sirbu and Cortez, 2013). Although the detailed mechanism of how ATR and ATM coordinately function during oogenesis in *Drosophila* is still unclear, it seems that both proteins have distinct functions. Specifically, ATM is primarily involved in DNA damage repair, whereas ATR is also involved in cell-cycle checkpoint regulation (Joyce et al., 2011).

Due to the fundamental role that *γ*H2AX (or *γ*H2Av) has in all DNA damage repair pathways, the recognition of this modification has become a standard assay for DSB and DNA damage detection. In this study, we find a strong accumulation of *γ*H2Av in mutant nurse cells, and reveal that mutation of the checkpoint gate keeper *ATR*/*mei-41* rescues the spindle-like phenotype in *su(Hw)* mutants. Previous observations have shown that an elevated frequency of DSBs is generally produced by mutations in DNA repair genes or by retrotransposon activity induced by mutations in components of the pi-RNA pathway. Both phenomena can lead to DNA damage and the activation of checkpoint repair pathways mediated by ATR and Chk2, which are phenotypically characterized by microtubule disorganization and translational repression of Grk (Chen et al., 2007; Klattenhoff et al., 2007; Mohn et al., 2014; Sienski et al., 2012). Our data, however, shows that *su(Hw)* mutant egg chambers display microtubule disorganization and Grk mislocalization, and that mutants for ATR and Chk1 *(grp)*, but not Chk2 *(mnk, lok*), partially rescue this phenotype. This important difference suggests that an alternative pathway, mediated by ATR and Chk1 is activated by a response to accumulation of DNA damage in *su(Hw)* mutant ovaries (Figure 10). This conclusion is further supported by our observation that DNA damage in *su(Hw)* mutant ovaries results from a pathway independent from transposable element activity.

A previous study reports that cutting the gene dosage of an RNA-binding protein, Rbp9, in half results in the rescue of female fertility in *su(Hw)*^*2/v*^ mutants (Soshnev et al., 2013). This study suggests that female sterility in *su(Hw)*^*2/v*^ mutants is due to the derepression of neuronal genes such as *rbp9* in the germline that are under the transcriptional control of Su(Hw) binding sites. Our results support the idea that Su(Hw) has roles during oogenesis that extend beyond transcriptional regulation. One possibility against this notion is that Su(Hw) may control the expression of genes involved in DNA damage signaling in the ovaries. However, published reports show no indication that Su(Hw) specifically controls the expression of genes involved in DNA repair or replication pathways (Baxley et al., 2011; Hsu et al., 2015). Instead, we suggest that null mutations of *su(Hw)* result in a complex multifactorial oogenesis phenotype. Sterility is caused mostly by the over-expression of neuronal genes such as *rbp9*, due to the lack of transcriptional repression activity of Su(Hw), whereas the spindle-like phenotype described here would be dependent on the structural activity of Su(Hw) in the genome through its insulator function. Thus, a reduction of the levels of *rbp9* is enough to restore oogenesis development but not sufficient to eliminate the ventralization phenotype induced by replication stress and the activation of the *ATR*/*chk1* checkpoint. Likewise, expression of *su(Hw)*::*eGFP* driven by *nanos*-*GAL4* can partially rescue fertility but does not completely eliminate the spindle-like phenotype, producing frequent mislocalization of Grk and ventralized eggshells.

DNA breaks in *su(Hw)* mutant females may be produced by two different mechanisms: an overall increase in the production of DNA breaks or a failure in the process of repairing DNA breaks. Intrinsic causes of DSBs in *Drosophila* female germline development are normally attributed to meiotic recombination, transposable element mobilization, or abnormal DNA replication. In this work, we have shown that excess of DNA damage in *su(Hw)* mutant ovaries does not result from abnormal meiotic recombination (Figure 4), or retrotransposon mobilization (Figure 5), suggesting that accumulation of DNA damage may result from abnormal genome replication during oogenesis. In ovaries, endoreplication is a specialized genome replication process that takes place within a limited time during oogenesis and produces polyploid nurse cells that supply nutrients for oocyte development. Cells undergoing endoreplication only pass through a G phase and a S phase, but do not undergo mitosis (M) (Lee et al., 2009). Euchromatin is duplicated early during endoreplication while heterochromatic sequences are duplicated at late S phase. Because S phase is shorter in each endoreplication cycle, some heterochromatic regions frequently lose the opportunity for replication. This loss of replication is called underreplication, and damaged DNA has been observed at the junction between replicated euchromatin and underreplicated heterochromatin regions (Hammond and Laird, 1985; Lilly and Spradling, 1996b; Yarosh and Spradling, 2014). In salivary gland polytene chromosomes, for example, the accumulated *γ*H2Av signals are detected at local underreplication sites (Andreyeva et al., 2008).

Because Su(Hw) has been associated with heterochromatin-euchromatin boundaries (Khoroshko et al., 2016), one possibility is that the absence of Su(Hw) in mutants disrupts the endoreplication process and leads to an excess of DNA damage at these sites. DNA breaks may result from stalled replication forks at the boundaries that fail to reinitiate replication (Lilly and Spradling, 1996a; Mirkin and Mirkin, 2007; Peng and Karpen, 2008). Interestingly, this mechanism has been suggested to explain the five lobes nuclear organization phenotype, which is typical in nurse cells during stages 1 to 5 of oogenesis, but that extends beyond stage 5 in *su(Hw)* and in other oogenesis mutants (Dej and Spradling, 1999). The 5 lobes structure is the result of homolog chromosome pairing during re-replication, and in normal nurse cells, these chromosomes separate and disperse in the nucleus after stage 5. The 5 lobes anomalous phenotype at later stages results from failure of chromosomes to disperse, which is likely due to unfinished replication forks that prevent sister chromatids from dispersing after stage 5 (Dej and Spradling, 1999).

Whereas research on insulator proteins has focused mostly on their roles in transcription, the possibility that chromatin insulators might also be involved in genome replication emerged recently after analysis of the genomic distributions of insulators and origin of replication proteins. This analysis revealed that a number of ORC and MCM2-7 helicase complex binding sites overlap with binding sites of Su(Hw). In addition, Su(Hw) is also capable of altering chromatin accessibility by recruiting the histone acetyltransferase SAGA and the BRAHMA chromatin remodeling complex, thereby creating a platform of low nucleosome density levels favorable for the recruitment of ORC and replication firing (Lu and Tower, 1997; Vorobyeva et al., 2013). In line with these findings suggesting a role for Su(Hw) in the regulation of DNA replication, we have also found a significant accumulation of H4K20me1 in *su(Hw)* mutant ovaries. H4K20me1 is highly regulated during the cell cycle and plays an important role in the licensing of origins of replication. PR-Set7 is the histone methyltransferase responsible for the monomethylation of H4K20 (H4K20me1). Mutations in PR-Set7 in mice models suggest that this protein has a role in DNA replication (Abbas et al., 2010; Tardat et al., 2010a). Specifically, mutations in *set7* that prevent PR-Set7 degradation during the cell cycle are lethal due to uncontrolled re-replication of the genome (Centore et al., 2010), and loss of PR-Set7 enzymatic activity also causes defects in origin of replication firing (Jorgensen et al., 2007; Tardat et al., 2007). Further experimental evidence has shown that the role of PR-Set7 in origins of replication depends on its specific mono-methyltransferase activity on H4K20. H4K20me1 functions as a substrate for subsequent methylation by Suv4-20h1 and Suv4-20h2 methyltransferases, which further dimethylate and trimethylate H4K20me1, respectively. H4K20me2 and H4K20me3 directly bind ORC and promote firing of replication forks at replication origins (Beck et al., 2012a).

Our finding showing increased levels of H4K20me1 in ovaries is reminiscent of mutations that prevent PR-Set7 degradation, increasing the frequency of H4K20me1 and leading to re-replication. The observation that Chk1, and not Chk2, is the kinase activated by ATR in response to replication stress, suggests that at least two checkpoint pathways (ATR/Chk1 and ATR/Chk2) can independently trigger microtubule disorganization and dorsoventral transformations (Figure 10). Convergence of the two pathways is most likely explained because of the functional role that DNA repair proteins, including ATR, Chk1, and Chk2, have in the MTOC, coordinating the cell cycle with DNA repair functions (Golan et al., 2010; Katsura et al., 2009; Shimada and Komatsu, 2009).

Altogether, our data suggest that DNA damage and genome instability may arise upon loss of Su(Hw) function in both germline and replicating somatic cells. Whether the DNA damage in *su(Hw)* mutants results from defects in genome organization that lead to replication stress, defects in the DNA repair pathway, or both is still unknown. However, our observation of chromosomal aberrations in dividing neuroblasts in *su(Hw)* mutants suggest this phenomenon expands also to replicating somatic cells and is not limited to the germline or to polytene chromosomes. Our study opens a new avenue to further understand the role of architectural proteins in genome function and genome stability.

## Materials and methods

### Fly genetics

All fly stocks were cultured on cornmeal-agar food with yeast at 25°C. The fly stocks used in this study are: *y*^*2*^*wct*^*6*^; *su(Hw)* ^*v*^*/TM6B*, a gift from Victor Corces (Emory University); *mei-41*^*D5*^, and w*; P{*GAL4*-*nos*.NGT40} (BDSC: 4442), which we refer to as *nos*-*GAL4* through the text, were gifts from Laura Lee (Vanderbilt University). *mei-w68* and P{*nanos*-*GAL4*::*VP16*}, which we refer to as *nanos*-*GAL4* throughout the text, were gifts from Bruce McKee (University of Tennessee). *mnk*^*6006*^, was a gift from Bill Theurkauff (UMass Worcester). *grpz*^*5170*^ *lok*^*30*^ and *grp*^*209*^ *lok*^*30*^ lines were a gift from Eric Weischaus (Princeton University). We also used Su(Hw)::*eGFP*[*yw*; P{*su(Hw)*::*eGFP*,w^+^}] and *w*^*1118*^;PBac(RB)*su(Hw)* ^*e04061*^/TM6B (BDSC: 18224).

### Immunofluorescence staining of ovaries

Three to five-day-old female ovaries were collected for ovary whole mount immunostaining as described previously (Page and Hawley, 2001). Briefly, tissues were fixed in 4% paraformaldehyde in 1:1 PBS and heptane (Sigma) and washed with PBST. Fixed tissues were incubated with blocking solution. Primary antibodies used for staining were as follows: FITC-conjugated mouse anti-tubulin (Sigma, 1:500), mouse anti-C(3)G (from Scott Hawley, Stowers Institute for medical research), rabbit anti-*eGFP* (Invitrogen, 1:100), rabbit anti-*γ*H2Av (Rockland, 1:5000), mouse anti-Orb, and anti-Grk (Developmental Studies Hybridoma Bank, 1:200). The following secondary antibodies were used at 1:200 dilution: FITC-conjugated anti-rabbit IgG, TexasRed-conjugated anti-rabbit IgG, and FITC-conjugated anti-mouse IgG (The Jackson Laboratory). F-actin staining was performed using Texas Red-X phalloidin (Life Technologies). Ovaries were stained with 4**′**, 6-diamidino-2-phenylindole (DAPI, 0.5 **μ**g/ml) and were mounted in Vectashield mounting medium (Vector Laboratories). Slides were analyzed under a Leica DM6000B wide-field fluorescence microscope equipped with a Hamamatsu ORCA-ER CCD camera and a HC PL FLUOTAR 20x/0.50NA objective. Image acquisition was performed using Simple PCI v6.6 (Hamamatsu Photonics). Images were processed using the AutoQuant 3D Deconvolution Algorithm utilizing an adaptive (blind) PSF implemented into Lecia Deblur (v2.3.2) software. Wildtype and mutant samples were prepared and imaged under identical conditions of immunostaining, microscope, camera, and software settings. Egg chamber stage was determined based on size (Sullivan et al., 2000) measured in FIJI (Schindelin et al., 2012).

### Documentation of eggshell phenotype

Two-to three-day-old *su(Hw)* mutant females carrying the *su(Hw)*::*eGFP* transgenes driven by P{*GAL4*-nos.NGT40} or P{*nanos*-*GAL4*::VP16}, were crossed to *yw* male flies. Eggs were collected for three days using grape juice agar plates containing wet yeast paste (Sullivan et al., 2000). The eggshell morphology was observed and documented using a Leica MZ16FQ stereomicroscope equipped with a Leica DFC420 digital camera.

### Western blot

Three to five-day-old female ovaries were dissected and homogenized in RIPA lysis buffer containing protease and phosphatase inhibitors (Roche). In some samples, stages 10 and older were removed manually under the dissecting microscope, previously to homogenization. After removal of stages 9 and older, wildtype and mutant ovaries contain the same developmental stages. Lysates were resolved on a 15% acrylamide gel, transferred overnight at 4°C, and probed with anti-*γ*H2Av (Rockland, 1:1000), rabbit anti-monomethylated H4K20 (Abcam, 1:1000) and mouse anti-lamin Dm0 (DSHB, 1:1000).

### Real-time PCR

Real-time PCR quantification of TE expression was carried out with ABGene (Rockford, IL) SYBR green PCR master mix. PCR conditions for each primer pair were tested to determine the efficiency of amplification and to ensure that the amplification was in the linear range. PCR products for each primer pair were amplified from cDNA using the BioRad iQ5 Multicolor Real-Time PCR detection system (Primers listed in Supporting Information Table 1). cDNA was reverse transcribed from at least three different RNA samples. Threshold Cycle (Ct) values were normalized to the Ct values of the housekeeping gene *rp49*. Change in expression level was calculated using the ΔΔCt method based on the Ct value for each PCR reaction (BioRad real-time PCR application guide). Results are presented as fold change in mutant relative to wildtype. The statistical significance of the results was calculated using Student’s t-tests.

### Chromosomal aberration (CAB) assay of larval brain squashes

Wandering third instar larvae were quickly dissected in PBS to obtain the brain. Brains were incubated in a small volume of 0.5% sodium citrate, pH 6.00, for 10 minutes. The brains were then fixed in 4% paraformaldehyde (2:1:1 acetic acid, 16% paraformaldehyde, dH_2_O) for 1 minute. The brains were squashed between a slide and a coverslip before staining with 4′, 6-diamidino-2-phenylindole (DAPI, 0.5 μg/mL) and were mounted in Vectashield mounting medium (Vector Laboratories). To determine the frequency of metaphasic cells that show chromosomal aberrations (CABs) in larval brains, pictures were taken of approximately fifty fields of view per sample that contained at least one metaphasic cell. Microscopy was performed as described above. Micrographs of neuroblast karyotypes were examined using FIJI (Schindelin et al., 2012) and each metaphasic cell was scored manually for the presence of CABs based on DAPI staining. Comparisons between genotypes were performed using two-tailed Mann-Whitney t-tests.

## Acknowledgements

We thank Dr. Laura Lee at Vanderbilt University for *P{GAL4-nos.NGT40}* and *mei-41*^*D5*^ stocks, Dr. Victor Corces at Emory University for *y*^*2*^*wct*^*6*^; *su(Hw)*^*v*^*/TM6B* stock, Dr. Bill Theurkauff at UMass Worcester for the *mnk*^*6006*^ stock, Dr. Eric Weischaus at Princeton University for the *grpz*^*5170*^ *lok*^*30*^ and *grp*^*209*^ *lok*^*30*^ lines, and Dr. Bruce McKee at the University of Tennessee for *P{nos-GAL4::VP16}* and *mei-w68* stocks. We appreciate the discussions provided by Dr. Bruce McKee and Dr. Albrecht von Arnim at the University of Tennessee, and a former Lab member Dr. Todd Schoborg for his support and several reagents. Also, we thank Tim Wesley for his help editing and making figures. The anti-Orb (4H8), lamin Dm0, Grk monoclonal antibodies, were developed by Drs. Paul Schedl, Paul A. Fisher, and Trudi Schupbach, respectively, and were obtained from the Developmental Studies Hybridoma Bank, created by the NICHD of the NIH and maintained at The University of Iowa, Department of Biology, Iowa City, IA 52242.

---

**Figure S1. Quantification of ring canals in *su(Hw)*^*v*/*e04061*^ egg chambers with abnormal nurse cell number.** The number of ring canals in egg chambers with either <15 nurse cells (top) or >15 nurse cells was quantified. 33% of egg chambers with <15 nurse cells contained <4 ring canals, whereas 66% contained 4 ring canals. 100% of egg chambers with >15 nurse cells contained >4 ring canals.

**Figure S2. Staging of egg chambers for Grk localization experiments.** The length of stage 9 (S9) egg chambers was first measured in wildtype flies. Egg chambers used for localization of Grk in mutant females is not significantly different from S9 egg chambers in wildtype. Egg chamber length was measured in arbitrary units. Comparisons between genotypes were performed using two-tailed Mann-Whitney t-tests.

