## Supplemental figures and table for "Mutations in the Insulator Protein Suppressor of Hairy Wing Induce Genome Instability"

| <15 Nurse Cells | >4 Ring Canals | <4 Ring Canals | 4 Ring Canals |
|---|---|---|---|
| 6 | 0 | 33.33% | 66.67% |
| >15 Nurse Cells | >4 Ring Canals | <4 Ring Canals | 4 Ring Canals |
| 2 | 100.00% | 0 | 0 |

**Figure S1**. Hsu et al.

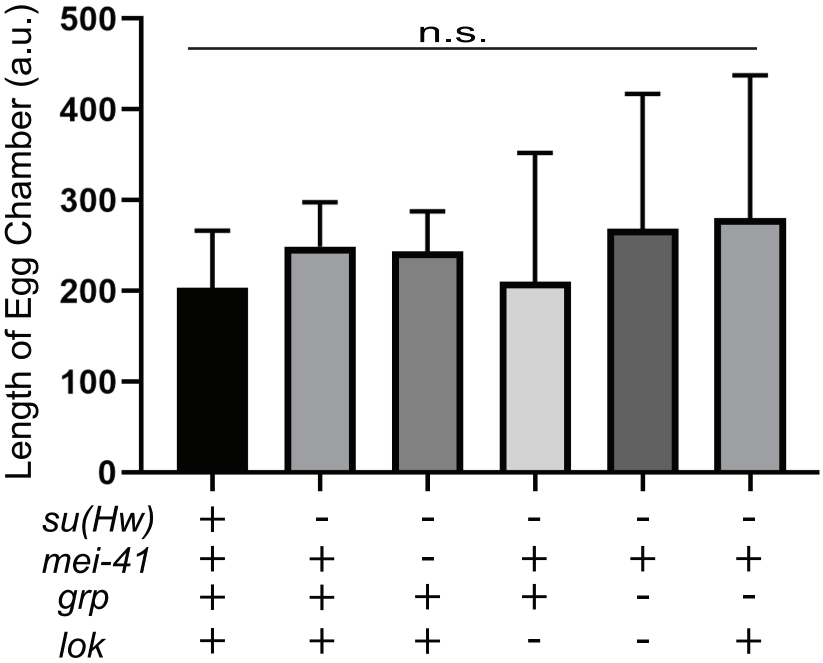


**Figure S2**. Hsu et al.

**Supplementary Table 1:** Transposable element primers

| **Transposable Elements** | **Forward primer (5'→3')** | **Reverse primer (5'→3')** | **Product size (bp)** |
|---|---|---|---|
| *Rp49* | TGTCCTTCCAGCTTCAAGATGACCATC | CTTGGGCTTGCGCCATTTGTG | 194 |
| *Gypsy* | TGGAAGCACCGCAAATCAAG | TCCAGGCCACATACTCGTC | 129 |
| *Jockey* | GCAGCACGGTACTCCTGAG | CAGGGTGCCAGACTCTGTC | 128 |
| *Doc3* | CTTCATGACCTTCATGCAAG | GCCATTAGCGTTCCAGGTA | 129 |
| *Nomad* | CAACGCCTCTCCAGTGTAC | GAGAAGGGTTTACGGACTGT | 143 |
| *X-element* | CCTTCGGCTACAGAACCTAG | GCAGCTTGATGACTGGTACTG | 134 |
| *RooA* | CAGAAGATGTTAACTCCAATTT | TCAATGAGTGTAGCTGTTTCG | 135 |
| *Baggins* | GGACTGTGTACCGATCGTG | GTGTTCAGCCAGTGCAGTG | 121 |
| *Beagle* | CTGACCATCAGCCTTTGAC | CAGAGCGTCGGCTACAGTA | 140 |
| *Stalker* | GTAGCAGACGCACTCTCAC | CCTAGGCAATAGTTCCTTG | 132 |
| *Gtwin* | ATGAAGTCACTCGGCAACCT | ACGCTTGGTAAAAGTATGCAATTG | 184 |
| *Tabor* | GGACCGACAACAAAGAAACATG | GAGAACTTTCGATACCTGAG | 123 |
| *412* | CCGTGTGATGGAATAATCGG | GGACAACTTGGGATCTTGCT | 181 |
| *Idefix* | GTACGGTACTGATCAACTG | GAATACTACTTTCACGTAGATTC | 120 |
| *Copia* | CCCTATTTGAAGCCGTGAGA | GACATGAGGGGTTGTTTGCT | 135 |
| *I-Element* | GCTCTTTCACCTCAACCATC | GCTAGCCAATGTAGTCTCGT | 140 |
